# O-GlcNAcylation regulates neurofilament-light assembly and function and is perturbed by Charcot-Marie-Tooth disease mutations

**DOI:** 10.1101/2023.02.22.529563

**Authors:** Duc T. Huynh, Jimin Hu, Jordan R. Schneider, Kalina N. Tsolova, Erik J. Soderblom, Abigail J. Watson, Jen-Tsan Chi, Chantell S. Evans, Michael Boyce

## Abstract

The neurofilament (NF) cytoskeleton is critical for neuronal morphology and function. In particular, the neurofilament-light (NF-L) subunit is required for NF assembly *in vivo* and is mutated in subtypes of Charcot-Marie-Tooth (CMT) disease. NFs are highly dynamic, and the regulation of NF assembly state is incompletely understood. Here, we demonstrate that human NF-L is modified in a nutrient-sensitive manner by O-linked-β-*N*-acetylglucosamine (O-GlcNAc), a ubiquitous form of intracellular glycosylation. We identify five NF-L O-GlcNAc sites and show that they regulate NF assembly state. Interestingly, NF-L engages in O-GlcNAc-mediated protein-protein interactions with itself and with the NF component α-internexin, implying that O-GlcNAc is a general regulator of NF architecture. We further show that NF-L O-GlcNAcylation is required for normal organelle trafficking in primary neurons, underlining its functional significance. Finally, several CMT-causative NF-L mutants exhibit perturbed O-GlcNAc levels and resist the effects of O-GlcNAcylation on NF assembly state, indicating a potential link between dysregulated O-GlcNAcylation and pathological NF aggregation. Our results demonstrate that site-specific glycosylation regulates NF-L assembly and function, and aberrant NF O-GlcNAcylation may contribute to CMT and other neurodegenerative disorders.

## Introduction

The unique morphology, homeostasis, and functions of neurons depend on a dynamic and highly regulated cytoskeleton, comprising actin, microtubule, and intermediate filament (IF) compartments^1^. Neurofilaments (NFs) are neuronal IFs, composed of three major “triplet” subunits (light, medium, and heavy) that form heterotypic polymers with each other and with two additional NF proteins, α-internexin (INA) and peripherin, in the central and peripheral nervous systems, respectively^2-4^. All NF proteins contain an α-helical coiled-coil rod domain flanked by amino-terminal head and carboxyl-terminal tail domains of varying lengths^2,3,5^. NF proteins assemble in discrete states, with two monomers first forming coiled-coils in the rod domain to create head-to-head dimers^2,3,5^. Two dimers then assemble in antiparallel fashion, forming a nonpolar tetramer^2,3,5^. Eight tetramers anneal laterally to form ∼65 nm unit-length filaments, which elongate end-to-end into short filaments and then compact radially into mature, fully assembled NFs of ∼10 nm diameter^2,3,5^. Due to their nonpolar nature, IFs cannot serve as tracks for molecular motors but instead exhibit viscoelastic properties distinct from the actin or microtubule networks^1,5-8^. For example, IFs are flexible under low strain but rigidify and resist breakage under applied force, with fully assembled filaments stiffening more than lower-order oligomers^1,5-12^. Therefore, the assembly state of all IFs, including NFs, is critical for their contributions to cell physiology.

Of the triplet proteins, NF-light (NF-L) is required for the structural integrity of axons^13-16^, has been detected at post-synaptic sites^17^, and directly interacts with the *N*-methyl-D-aspartate receptor to influence high-order brain functions^18-20^. Ablating the *NEFL* gene, which encodes NF-L, impairs maturation of regenerating myelinated axons^13^, dendritic arborization^14^, and peripheral nerve regeneration in mice^15^. Motor neurons derived from human *NEFL*^*-/-*^ induced pluripotent stem cells (iPSC) show reduced axonal caliber, dysregulated mitochondrial motility, and decreased electrophysiological activity^16^. Moreover, NF-L assembly state and functions are perturbed in a range of nervous system disorders. *NEFL* mutations cause some subtypes of Charcot-Marie-Tooth (CMT) disease^21,22^, an inherited peripheral neuropathy characterized by progressive atrophy of the distal limb muscles that leads to sensory loss and tendon reflex defects^23^. CMT-causative mutations in *NEFL* or other genes result in NF aggregation and aberrant motility of neuronal mitochondria^24-30^, underlining the physiological importance of NF-L function. Aggregation of wild type (WT) NF-L and other NF proteins is also a pathological hallmark of a variety of neurological conditions, including Alzheimer’s disease (AD)^31^, Parkinson’s disease (PD)^32^, amyotrophic lateral sclerosis (ALS)^33^, giant axonal neuropathy^25^, and spinal muscular atrophy^34^. Recently, NF-L levels in the cerebrospinal fluid (CSF) and blood have emerged as powerful biomarkers of neuronal injury and nervous system disorders^3,4,35^, showing promise for early diagnosis in a variety of clinical settings^36-40^. Given this broad pathophysiological significance, elucidating the regulation of NF assembly state and functions is a key goal. However, the molecular mechanisms governing NFs remain incompletely understood.

One major mode of NF regulation is likely through various post-translational modifications (PTMs)^2,3,41-46^. Prior proteomic studies indicated that rodent NFs are modified by O-linked-β-*N*-acetylglucosamine (O-GlcNAc)^47-52^, an abundant intracellular form of glycosylation reversibly decorating serine and threonine sidechains on many nuclear, cytoplasmic, and mitochondrial proteins (Figure 1A)^53-55^. In mammals, O-GlcNAc is added by O-GlcNAc transferase (OGT) and removed by O-GlcNAcase (OGA), both ubiquitous nucleocytoplasmic enzymes^53-55^. O-GlcNAc cycling is essential, as deletion of *OGT* or *OGA* is lethal in mice^56,57^. O-GlcNAc is common in many mammalian tissue types^53-55^ and is especially prevalent in the brain and post-synaptic densities^58,59^. Ablating the *OGT* gene in specific populations of murine dopaminergic^51^ or hypothalamic^60,61^ neurons causes cellular and behavioral defects, demonstrating the importance of O-GlcNAc to brain function. At the molecular level, O-GlcNAc mediates various aspects of mammalian neuronal biology^62,63^, and O-GlcNAcylation of disease-relevant substrates, such as tau^64^ or α-synuclein^65^, is dysregulated in multiple clinically important neurological disorders^66,67^. Manipulating O-GlcNAc in the nervous system has shown therapeutic promise in recent pre-clinical studies and clinical trials alike^68-75^. For example, multiple reports showed that elevating O−GlcNAcylation via treatment with the small molecule OGA inhibitor Thiamet-G^69^ reduced proteotoxicity, cognitive deficits, and behavioral dysfunction in AD rodent models^70,71^. Based on these and other promising pre-clinical data^68,72^, at least three OGA inhibitors (MK-8719, LY3372689, ASN90) have entered human clinical trials^73-75^, paving the way for future pharmacological modulation of O-GlcNAc on substrates, such as NF proteins, in human patients. However, despite its potential clinical significance, the functional impact of O-GlcNAcylation on human NF proteins has never been studied systematically.

**Figure 1:**
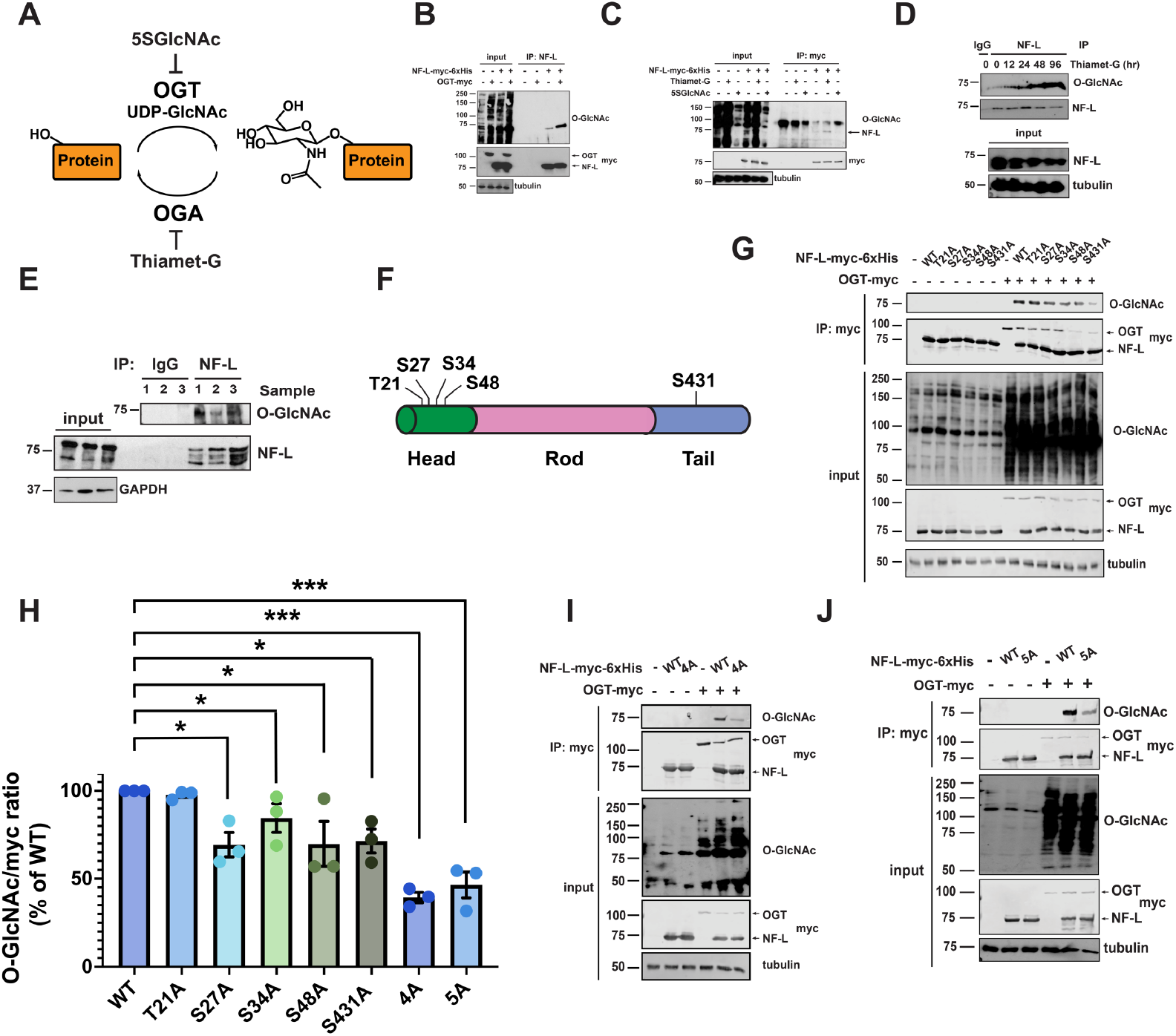
Site-specific O-GlcNAcylation of the human NF-L head and tail domains. (A) O-GlcNAc transferase (OGT) uses the nucleotide-sugar UDP-GlcNAc to add O-GlcNAc to serine or threonine residues of intracellular proteins, and O-GlcNAcase (OGA) catalyzes its removal. OGT and OGA can be inhibited by the small molecules 5SGlcNAc and Thiamet-G, respectively. (B) 293T cells were transfected with NF-L-myc-6xHis ± OGT-myc for 24 hrs, and lysates were analyzed by NF-L IP and IB. (C) 293T cells were transfected with NF-L-myc-6xHis for 24 hrs, treated with 50 µM Thiamet-G or 50 µM 5SGlcNAc for 30 hrs, and lysates were analyzed by myc IP and IB. (D) SH-SY5Y cells were cultured in 50 µM Thiamet-G for the times indicated, and lysates were analyzed by control (IgG) or NF-L IP and IB. (E) Human temporal cortex homogenates from three donors (1-3) were analyzed by control (IgG) or NF-L IP and IB. (F) Five O-GlcNAc sites identified by MS in this and prior studies are indicated on the NF-L domain structure. (G) *NEFL*^-/-^ 293T cells were transfected with NF-L-myc-6xHis WT or single-point glycosite mutants ± OGT for 24 hrs, and lysates were analyzed by myc IP and IB. (H) Normalized O-GlcNAc signal (O-GlcNAc/myc ratio) was calculated for the experiments performed in (G, I-J) (n=3). (I, J) *NEFL*^-/-^ 293T cells were transfected with NF-L-myc-6xHis WT or NF-L^4A^ (I) or NF-L^5A^ (J) ± OGT for 24 hrs, and lysates were analyzed by myc IP and IB. Throughout: n.s., not significant; **, p < 0.005; ***, p < 0.0005.

Here, we demonstrate that human NF-L is O-GlcNAc-modified in cell culture models, primary neurons, and post-mortem brain tissue. NF-L assembly state and function are regulated by site-specific O-GlcNAcylation, as elevating O-GlcNAc drives NF-L to lower-order oligomeric states, reduces the prevalence of full-length filaments, and alters organelle motility in primary hippocampal neurons. At the molecular level, we show that NF-L O-GlcNAc is nutrient-responsive and mediates both homotypic NF-L/NF-L and heterotypic NF-L/INA interactions, revealing previously unknown, glycan-mediated interactions among NF components. Further, we observed aberrant O-GlcNAc levels on CMT-causative NF-L mutants and hypoglycosylation of NF-L when CMT mutations lie proximal to glycosites. Interestingly, hypoglycosylated CMT NF-L mutants formed aggregates that were insensitive to the normal assembly state effects of O-GlcNAcylation, compared to WT NF-L. Together, our results indicate that site-specific O-GlcNAcylation is an important mode of NF regulation and may be dysregulated in neurological disorders.

## Results

### Site-specific O-GlcNAcylation of the human NF-L head and tail domains

Several prior studies reported the O-GlcNAcylation of rodent NF proteins^47-52^. However, the existence and functions of O-GlcNAc on human NFs had not been examined systematically. To address this knowledge gap, we first focused on NF-L because it is essential for NF assembly *in vivo*^2-4^ and is mutated in subtypes of CMT^21,22^. In cells expressing tagged human NF-L, OGT co-expression induced global O-GlcNAcylation and elevated NF-L O-GlcNAc levels (Figure 1B). In contrast, expression of OGT^H498A^, a glycosyltransferase-dead mutant^76^, did not affect NF-L O-GlcNAcylation, confirming the specificity of the assay (Figure s1A). Inhibiting OGA or OGT with the small molecule Thiamet-G or peracetylated 5-thio-GlcNAc (5SGlcNAc)^77^ elevated or reduced NF-L O-GlcNAcylation, respectively (Figure 1C). In human neuroblastoma SH-SY5Y cells, Thiamet-G treatment increased endogenous NF-L O-GlcNAcylation in a time-dependent manner (Figure 1D). Furthermore, we detected endogenous NF-L O-GlcNAcylation in post-mortem human temporal cortex, frontal cortex, and parietal cortex tissue samples (Figures 1E, s1B). Mild β-elimination of O-linked glycans^78^ extinguished the signal on these anti-O-GlcNAc immunoblots (IBs), confirming it as authentic (Figure s1C). These results demonstrate that human NF-L is dynamically O-GlcNAcylated in culture models and *in vivo*.

Previous reports identified several O-GlcNAcylation sites on rodent NF-L orthologs^47-52^, but directed studies of human NF glycosites were lacking. To discover human NF-L glycosites, we epitope-tagged the *NEFL* genomic locus of human cells via established CRISPR/Cas9 methods^79^ (Figure s1D) and affinity-purified endogenous NF-L from cultures treated with Thiamet-G, a standard tactic to improve the technically challenging detection of O-GlcNAc moieties^80-83^. Mass spectrometry (MS) analysis identified two novel glycosites (S48, S431), complementing the previously reported glycosites from rodent NF-L^47,48^ (T21, S27, S34 – human numbering of cognate rodent residues) (Figure 1F). Next, we created an S/T→A mutation at each of these residues and measured the effects on total NF-L O-GlcNAcylation by immunoprecipitation (IP) and quantitative fluorescent IB to determine major glycosites. While all NF-L constructs displayed comparable basal levels of O-GlcNAcylation (Figure s1E), each mutant except T21A exhibited lower OGT-induced O-GlcNAcylation than WT (Figure 1G-H), suggesting that NF-L could be simultaneously O-GlcNAcylated at multiple residues in response to upstream stimuli. Consistent with this hypothesis, an NF-L mutant with all four head-domain glycosites changed to alanine (NF-L^4A^) exhibited significantly less induced O-GlcNAcylation, compared to WT (Figure 1H-I). The single S431A mutation in the tail domain glycosite also dramatically reduced NF-L O-GlcNAcylation, whereas a compound mutant lacking the four head and one tail domain sites (NF-L^5A^) exhibited total O-GlcNAcylation similar to the NF-L^4A^ mutant (Figure 1H and J), suggesting potential crosstalk between the head and tail domains during OGT modification. These data demonstrate the inducible, site-specific O-GlcNAcylation of human NF-L.

### NF-L O-GlcNAcylation influences assembly state and filament formation

The NF-L head domain is required for its assembly^2-4^, and other PTMs, such as phosphorylation, on distinct head domain residues regulate this process^42,43,46^. Therefore, we examined whether head domain O-GlcNAcylation affects NF-L assembly state. We used an established differential extraction assay that biochemically separates distinct IF assembly states (Figure 2A)^81,84^ to determine the effects of O-GlcNAcylation on the NF-L network. As expected, WT NF-L extracted overwhelmingly into a denaturing urea buffer, indicating that it was assembled into full-length filaments, which are poorly soluble in non-denaturing buffers (Figure 2B-C)^81,84^. In multiple human cell types, OGT co-expression significantly increased the solubility of NF-L in non-denaturing buffers of low or high ionic strength (Figures 2B-C and s2A-B), indicating that increased global O-GlcNAcylation drives NF-L to lower-order assembly states and reduces the prevalence of full-length filaments. In contrast, co-expression of OGT^H498A^ did not affect the NF-L extraction profile (Figure s2C), demonstrating a requirement for glycosyltransferase activity. To confirm these results via an independent approach, we turned to immunofluorescence assays (IFA). We re-expressed NF-L in *NEFL*^-/-^ SH-SY5Y cells (Figure 2D), with or without OGT co-expression. Consistent with our differential extraction data, IFA showed that NF-L formed intact filaments when expressed alone, but both filaments and lower-order oligomeric states were observed upon OGT co-expression (Figure 2D). These results demonstrate that increased global O-GlcNAc levels drive NF-L from a full-length filament assembly state to lower-order states.

**Figure 2:**
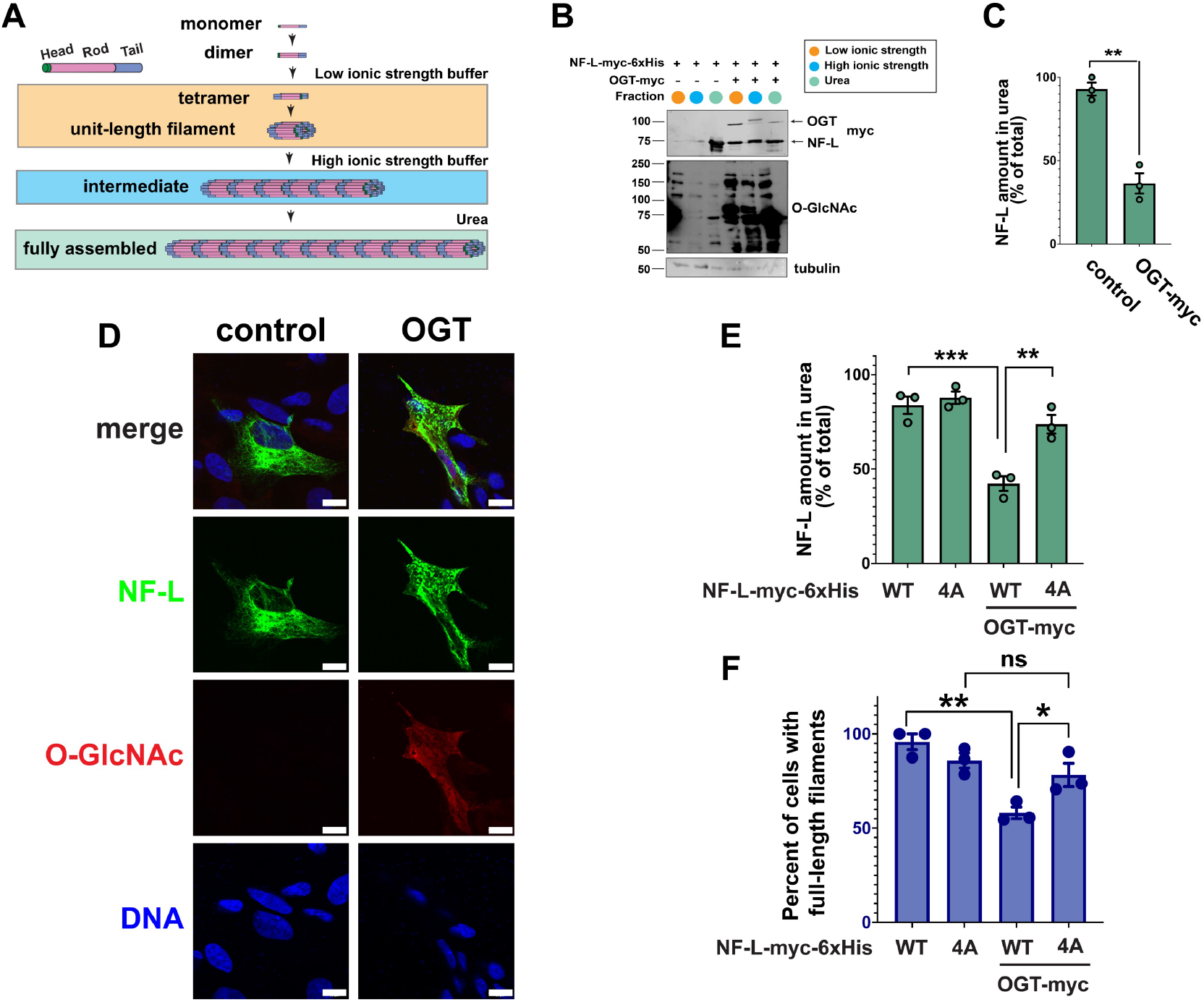
NF-L O-GlcNAcylation influences NF-L assembly state and filament formation. (A) Differential extraction assay separates discrete assembly states of NF-L. Low ionic strength, low-order assembly states; high ionic strength, intermediate assembly states; 8 M urea, fully assembled NFs. (B) *NEFL*^-/-^ 293T cells were transfected with NF-L-myc-6xHis ± OGT for 24 hrs and analyzed by differential extraction and IB. (C) NF-L amount extracted into urea buffer was calculated as percent of total NF-L across three fractions from the experiment described in (B) (n=3). (D) *NEFL*^-/-^ SH-SY5Y cells were transfected with NF-L-myc-6xHis ± OGT for 24 hrs and analyzed by IFA. Scale bar: 10 µm. (E) *NEFL*^-/-^ 293T cells were transfected with WT or NF-L^4A^-myc-6xHis ± OGT for 24 hrs and analyzed by differential extraction and IB. NF-L amount extracted into urea buffer was calculated as percent of total NF-L across three fractions (n=3). (F) *NEFL*^-/-^ SH-SY5Y cells were transfected with WT or NF-L^4A^-myc-6xHis ± OGT for 24 hrs and analyzed by IFA. Quantification of percent of cells with full-length NFs was performed by a blinded researcher (n=3). Throughout: *, p < 0.05; **, p < 0.005; ***, p < 0.0005.

OGT has thousands of substrates in human cells^53-55^ and could impact NF-L assembly state through direct and/or indirect mechanisms. To determine whether O-GlcNAcylation of NF-L itself governs NF assembly state, we performed differential extraction assays on WT and glycosite mutants of NF-L expressed in a tractable *NEFL*^-/-^ human cell system (Figure s2D-F). Consistent with our observation of uniform baseline glycosylation across NF-L WT and mutants (Figure s1E), the individual glycosite mutants displayed no changes in extraction profile, compared to WT, when expressed alone (Figure s2G-H). In contrast, in the presence of OGT co-expression, NF-L^4A^ resisted the shift to lower-order assembly states exhibited by the WT protein (Figure 2E), demonstrating that O-GlcNAcylation at specific sites in the NF-L head domain influences its assembly state. We obtained similar results in *NEFL*^-/-^ SH-SY5Y cells using an independent assay (IFA): At baseline, NF-L^4A^ displayed somewhat reduced prevalence of full-length filaments, compared to WT, but OGT expression did not decrease the NF-L^4A^ full-length filament population, as it does for WT (Figure 2F). Together, these results demonstrate that site-specific O-GlcNAcylation in the head domain regulates NF-L assembly state.

### NF-L head domain O-GlcNAcylation is required for organelle motility regulation

The NF network regulates organelle motility, and loss of NF-L causes accelerated mitochondrial movement in several neuronal model systems^16,24^. Therefore, we used live-cell fluorescence microscopy methods^85^ to test whether NF-L O-GlcNAcylation impacts organelle motility. Consistent with prior studies, expression of WT NF-L in cultured rat hippocampal neurons reduced mitochondrial total displacement, run length, and speed (Figure 3A-B). By contrast, expression of NF-L^4A^ did not significantly affect mitochondrial motility, as results from mock-transfected and NF-L^4A^-expressing neurons were statistically indistinguishable (Figure 3B). Notably, expression of WT NF-L increased lysosomal motility (displacement, run length, speed), whereas NF-L^4A^ expression decreased these parameters (Figure 3C-D). The NF network indirectly influences organelle motility by creating steric barriers and/or interactions with the microtubule cytoskeleton^86,87^, which forms the tracks for motor-driven organelle transport^88^. Expression of WT NF-L or NF-L^4A^ did not detectably affect the neuronal microtubule network (Figure s3), ruling out a general cytoskeletal disruption caused by loss of NF-L glycosylation. These results indicate that NF-L head domain O-GlcNAcylation modulates organelle motility in primary neurons.

**Figure 3:**
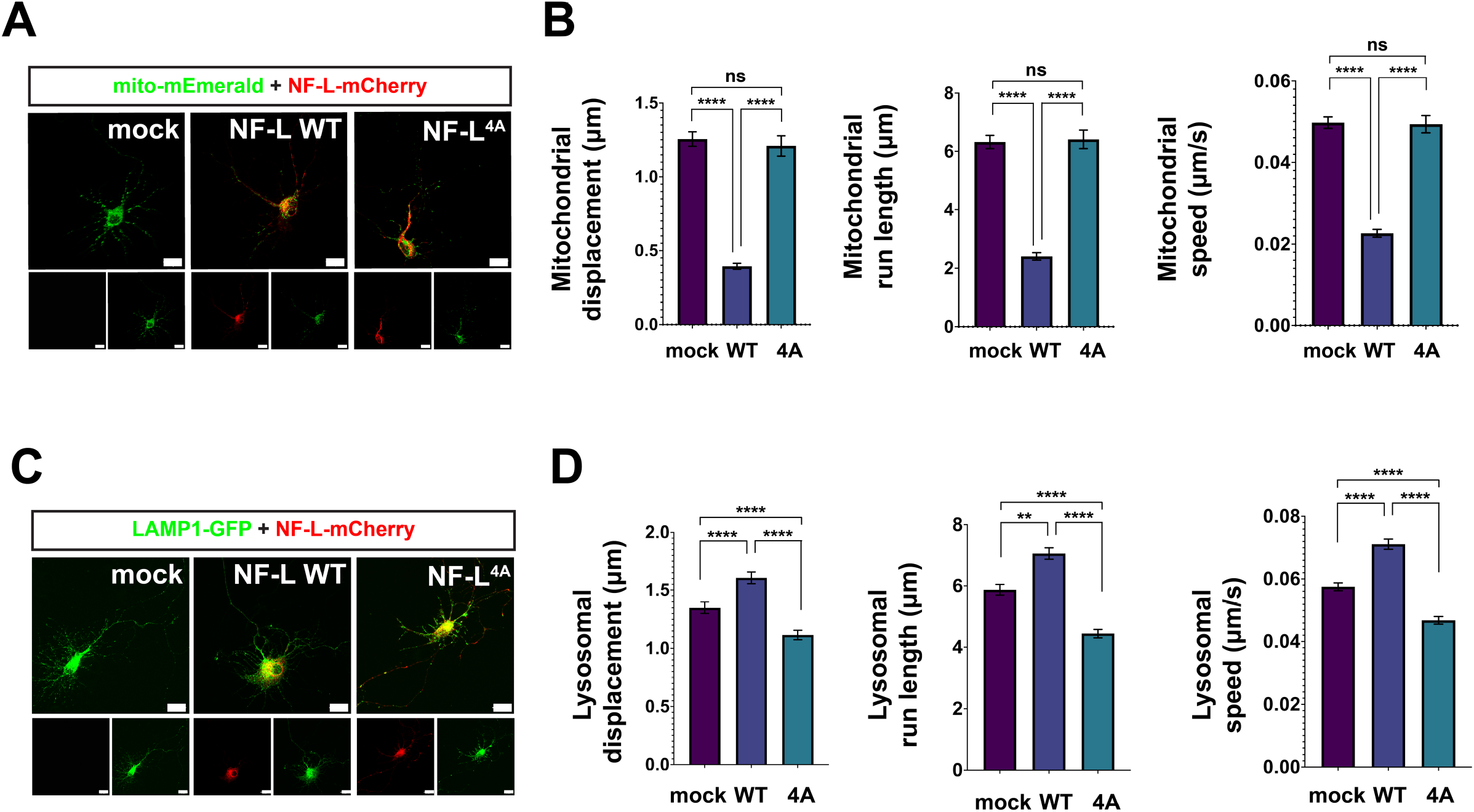
NF-L head domain O-GlcNAcylation is required for organelle motility regulation. (A) Cultured E18 rat hippocampal neurons at day 6 *in vitro* were transfected with WT or NF-L^4A^-mCherry + mito-mEmerald (mitochondrial marker) for 24 hrs and analyzed by live-cell fluorescence microscopy. Scale bar: 20 µm. (B) Mitochondrial motility in axons of neurons treated as in (A) was imaged every 5 s for 5 min and analyzed to obtain organelle displacement, run length, and speed. Data were assessed by Mann-Whitney test/Dunn’s post-hoc correction (n=3). (C) Cultured rat hippocampal neurons at day 6 *in vitro* were transfected with WT or NF-L^4A^-mCherry + LAMP1-GFP (lysosomal marker) for 24 hrs. Scale bar: 20 µm. (D) Lysosomal motility in axons of neurons treated as in (C) was imaged every 5 s for 5 min and analyzed to obtain displacement, run length, and speed. Data were analyzed by Mann-Whitney test/Dunn’s post-hoc correction (n=2). Throughout: n.s., not significant; **, p < 0.005; ****, p < 0.0001.

### Nutrient dependence of NF-L O-GlcNAcylation

Our organelle motility assays indicate that basal levels of O-GlcNAcylation are required for some NF-L functions because WT NF-L and NF-L^4A^ exhibited distinct phenotypes even in the absence of applied stimuli (Figure 3). However, elevating global O-GlcNAc levels with an experimental stimulus (e.g., OGT expression) revealed phenotypes that were not evident at baseline (Figures 2, s2), indicating that upstream signals might regulate NF-L by inducing or inhibiting its O-GlcNAcylation. One such candidate stimulus is a change in nutrient or growth factor availability. O-GlcNAc is well known to act in part as a sensor of nutrients, such as glucose and glutamine, which are biosynthetic precursors of uridine diphosphate (UDP)-GlcNAc, the nucleotide-sugar cofactor used by OGT (Figure 1A)^53-55^. Moreover, we have previously shown that fluctuations in nutrients or growth factors affect the O-GlcNAcylation of gigaxonin, a ubiquitin E3 ligase adaptor that targets NF proteins for proteasome-mediated destruction^82^. These results provided a precedent for metabolite-dependent regulation of NF-L through O-GlcNAc signaling. As a first step towards determining whether the O-GlcNAcylation of NF-L itself is affected by nutrient fluctuations, we starved cells of glucose, glutamine, or serum and quantified NF-L O-GlcNAcylation (Figure 4). Interestingly, all three starvation treatments resulted in significant reductions in NF-L glycosylation but not NF-L expression (Figure 4). These results establish nutrient and growth factor fluctuations as candidate stimuli that may impact NF-L O-GlcNAcylation and function *in vivo*.

**Figure 4:**
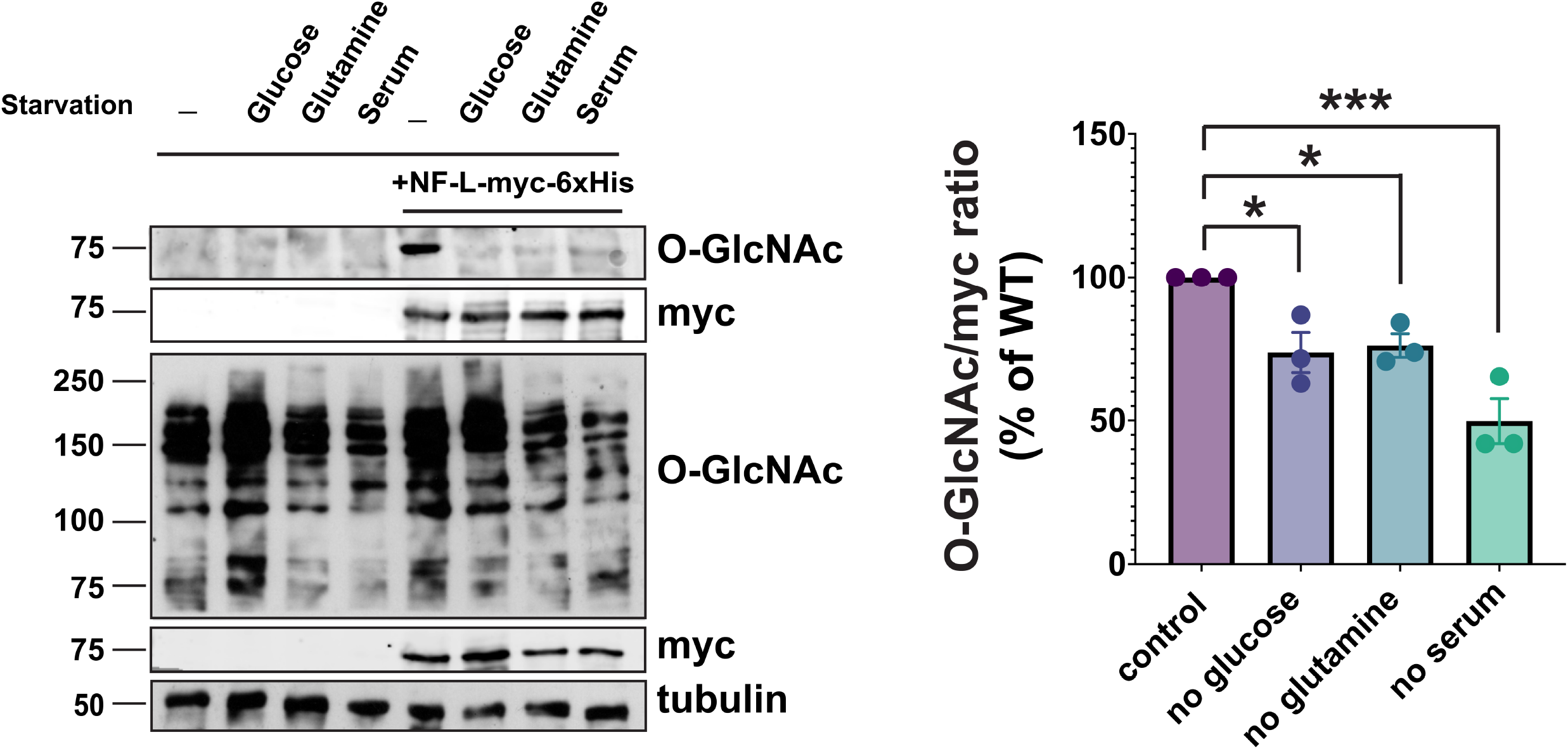
Nutrient dependence of NF-L O-GlcNAcylation. Left: *NEFL*^-/-^ 293T cells were transfected with NF-L-myc-6xHis for 24 hrs, grown in control medium or medium without glucose, glutamine, or serum for 48 hrs, and analyzed by myc IP and IB. Right: Normalized O-GlcNAc signal (O-GlcNAc/myc ratio) was calculated (n=3). *, p < 0.05; ***, p < 0.0005.

### Direct, O-GlcNAc-mediated interactions between NF-L and INA

We next sought to define the molecular mechanism by which O-GlcNAc influences NF-L assembly state. O-GlcNAc can mediate protein-protein interactions (PPIs) on a range of substrates^53-55,89^, and we have previously shown that the IF protein vimentin engages in homotypic, O-GlcNAc-mediated PPIs that are essential for filament formation^81^. Since NF-L both self-associates^90^ and co-polymerizes with other NF proteins into higher-order complexes^2^, we hypothesized that NF-L O-GlcNAcylation might influence NF assembly states by mediating PPIs. However, physiological O-GlcNAc-mediated interactions are often low-affinity and sub-stoichiometric, making them technically challenging to characterize^53-55,89^. To overcome this obstacle, we employed a chemical biology method to capture endogenous O-GlcNAc-mediated PPIs (Figure 5A)^91^. Briefly, live cells are first metabolically labeled with a precursor form of “GlcNDAz,” a GlcNAc analog bearing a diazirine photocrosslinking moiety^91^. GlcNDAz is metabolized to the nucleotide-sugar UDP-GlcNDAz, which is accepted by OGT, resulting in the installation of O-GlcNDAz moieties onto native substrates^91^. Brief UV treatment of GlcNDAz-labeled cells affords the covalent *in situ* crosslinking of O-GlcNDAz glycans to direct binding partner proteins within ∼2-4 Å of the sugar^91^. Therefore, GlcNDAz allows the capture, purification, and characterization of physiological O-GlcNAc-mediated PPIs^91^.

**Figure 5:**
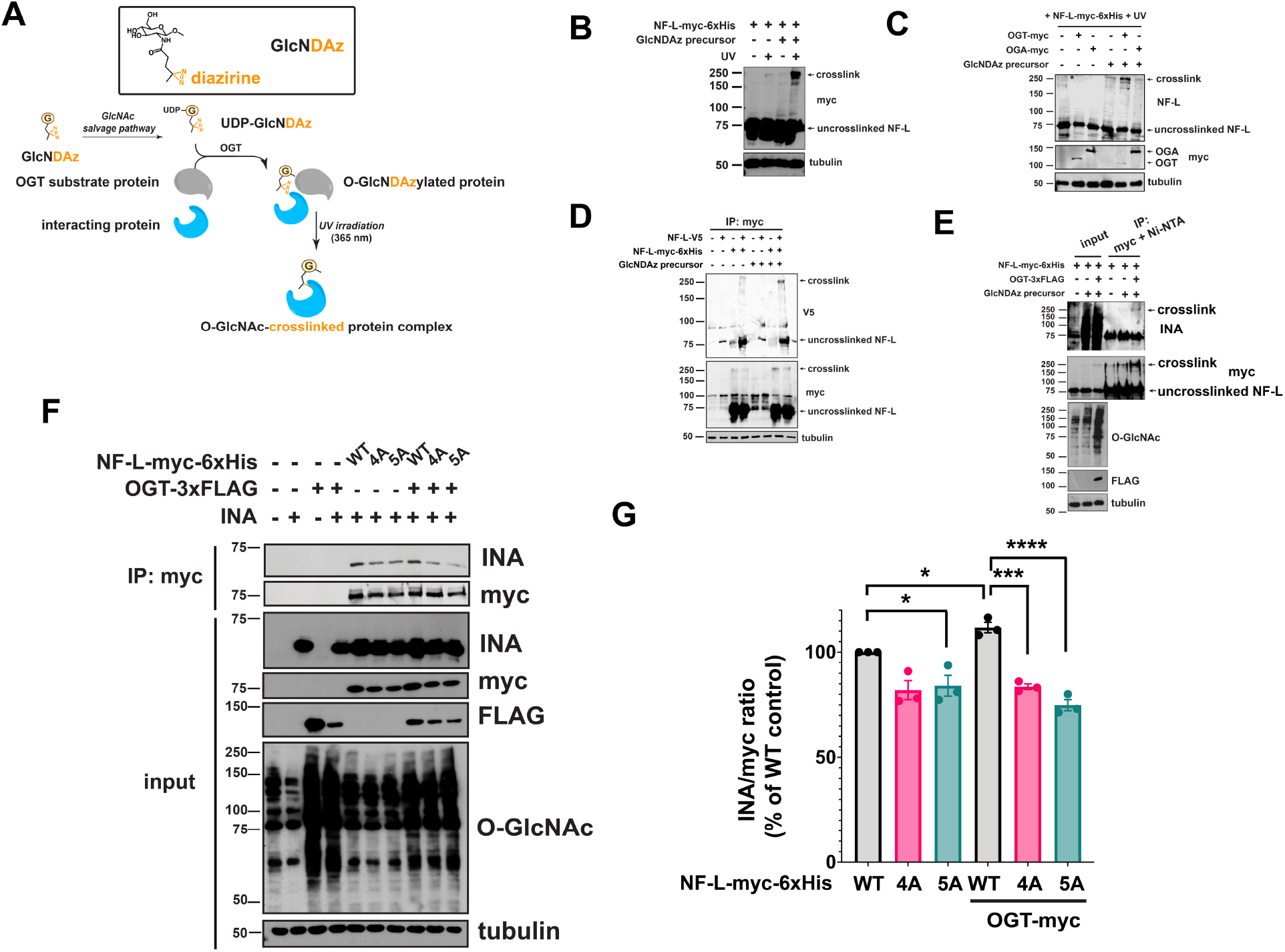
Direct, O-GlcNAc-mediated interactions between NF-L and INA. (A) GlcNDAz, a GlcNAc analog that bears a diazirine photocrosslinking moiety, can label OGT substrates in living cells. Brief UV treatment affords the covalent *in situ* crosslinking of O-GlcNDAz glycans to binding partner proteins, as described^91^. (B) 293T cells were transfected with NF-L-myc-6xHis ± 100 µM GlcNDAz for 48 hrs, subjected to UV crosslinking, and analyzed by IB. (C) 293T cells were transfected with NF-L-myc-6xHis ± OGT-myc or OGA-myc ± 100 µM GlcNDAz for 48 hrs, subjected to UV crosslinking, and analyzed by IB. (D) *NEFL*^-/-^ 293T cells were transfected with NF-L-myc-6xHis + NF-L-V5 ± 100 µM GlcNDAz for 48 hrs and analyzed by myc IP and IB. (E) *NEFL*^-/-^ 293T cells were transfected with NF-L-myc-6xHis ± OGT-3XFLAG ± 100 µM GlcNDAz for 48 hrs and analyzed by tandem myc IP/Ni-NTA purification and IB. (F) *NEFL*^-/-^ 293T cells were transfected with NF-L-myc-6xHis, NF-L^4A^-myc-6xHis or NF-L^5A^-myc-6xHis ± INA ± OGT-3XFLAG for 48 hrs and analyzed by myc IP and IB. (G) INA/myc ratio was calculated for the experiment described in (F) (n=3). *, p < 0.05; ***, p < 0.0005; ****, p < 0.0001.

We first performed GlcNDAz crosslinking on cells expressing WT NF-L. NF-L (∼62 kDa predicted molecular weight; ∼75 kDa on SDS-PAGE) crosslinked into ∼200-250 kDa complexes in a diazirine- and UV-dependent manner (Figure 5B). As expected, in the same samples, the heavily O-GlcNAcylated nucleoporin-62 also crosslinked, (positive control), whereas tubulin, which is not an OGT substrate, did not (negative control) (Figure s4A). Co-expression of OGT potentiated GlcNDAz crosslinking of NF-L, whereas OGA co-expression reduced it (Figure 5C), further supporting that the crosslinks are specific and are mediated by O-GlcNAcylation. These results indicate that NF-L engages in direct, O-GlcNAc-mediated PPIs.

We next identified the O-GlcNAc-mediated binding partners of NF-L. Because NF-L engages in both homotypic and heterotypic PPIs, we tested both types of interaction. First, we co-expressed two NF-L constructs with distinct epitope tags, performed GlcNDAz crosslinking, and analyzed the samples by IP/IB (Figure 5D). NF-L molecules with each tag were present in the same crosslinked complexes (Figure 5D), suggesting that NF-L engages in homotypic O-GlcNAc-mediated PPIs. Second, to identify other O-GlcNAc-mediated binding partners of NF-L, we affinity-purified crosslinked complexes from vehicle (DMSO)-treated and GlcNDAz-treated samples and analyzed them by MS. Interestingly, MS identified INA in the +GlcNDAz sample (186 INA peptides by spectral count) but not in the vehicle control (0 INA peptides) (Figure s4B). INA is a comparatively low-abundance NF protein that heteropolymerizes with the triplet proteins^92^ and aggregates in NF inclusion body disease, a form of frontotemporal dementia^93,94^. Tandem affinity purification experiments revealed that OGT expression increased the levels of both NF-L and INA in GlcNDAz-crosslinked complexes (Figure 5E), corroborating the MS results. Importantly, co-IP of WT NF-L and INA was potentiated by co-expression of OGT, confirming the O-GlcNAc-mediated interaction between these proteins in a GlcNDAz-independent assay (Figure 5F-G). Moreover, NF-L^4A^ and NF-L^5A^ exhibited reduced interaction with INA, compared to WT, and OGT co-expression failed to potentiate the interaction between INA and either NF mutant (Figure 5F-G). Taken together, these results demonstrate that O-GlcNAc moieties on NF-L engage in direct PPIs with other NF-L molecules and with INA, revealing previously unknown glycan-mediated interactions within the NF network.

### NF-L O-GlcNAcylation is dysregulated by CMT-causative mutations

CMT-causative *NEFL* mutations trigger NF-L accumulation and aggregation in neurons^26-30^, and WT NF protein aggregation is a feature of many neurodegenerative disorders^25,31-34^. Given our results demonstrating that O-GlcNAc influences NF-L assembly state and the many well-established connections among O-GlcNAcylation, protein aggregation, and neurodegeneration in general^62^, we tested whether NF dysfunction impacted NF-L O-GlcNAcylation. We generated NF-L constructs for 14 point-mutants that cause CMT^21,22^ and quantified their O-GlcNAcylation by IP/IB (Figure 6A-B). Strikingly, most (8/14) mutants displayed significantly altered O-GlcNAcylation, compared to WT, with six hypoglycosylated and two hyperglycosylated (Figure 6A-B). In particular, the four CMT mutations lying near NF-L glycosites (P8L, P22R, P22S, P440L) greatly reduced or completely abolished NF-L O-GlcNAcylation (Figure 6A-B). Because our results showed that NF-L O-GlcNAcylation modulates NF assembly state (Figure 2) and mediates PPIs (Figure 5), we hypothesized that hypoglycosylated CMT mutants would exhibit aberrant assembly. Consistent with literature reports demonstrating the formation of insoluble aggregates by CMT-causative NF-L mutants^30^, differential extraction experiments revealed that all four hypoglycosylated NF-L CMT mutants were predominantly insoluble under basal conditions (Figure 6C-D). Notably, elevating O-GlcNAc levels by OGT co-expression failed to drive these NF-L mutants into soluble assembly states, as it does for WT (Figure 6C-D). To confirm and extend these findings, we performed IFA experiments in SW13 vim^-^ cells, a tractable and well-established model system that lacks all cytoplasmic IFs, frequently used for imaging studies of IF protein mutations^95^. In line with prior reports and our own results, WT NF-L formed full-length, heteropolymeric filaments when co-expressed with INA, whereas most CMT mutants did not, instead forming punctate aggregates (Figure 6E-F). Interestingly, GlcNDAz experiments showed aberrant, enhanced crosslinking by aggregate-forming NF-L CMT mutants, compared to WT or to the filament-forming mutants (A149V, I384F, K443N, K467N) (Figure 6G). All together, these data demonstrate that many CMT-causative mutants exhibit abnormal NF-L O-GlcNAcylation, PPIs, and assembly states. Based on our results, we propose a working model for the function of NF-L O-GlcNAcylation (Figure 7), discussed below.

**Figure 6:**
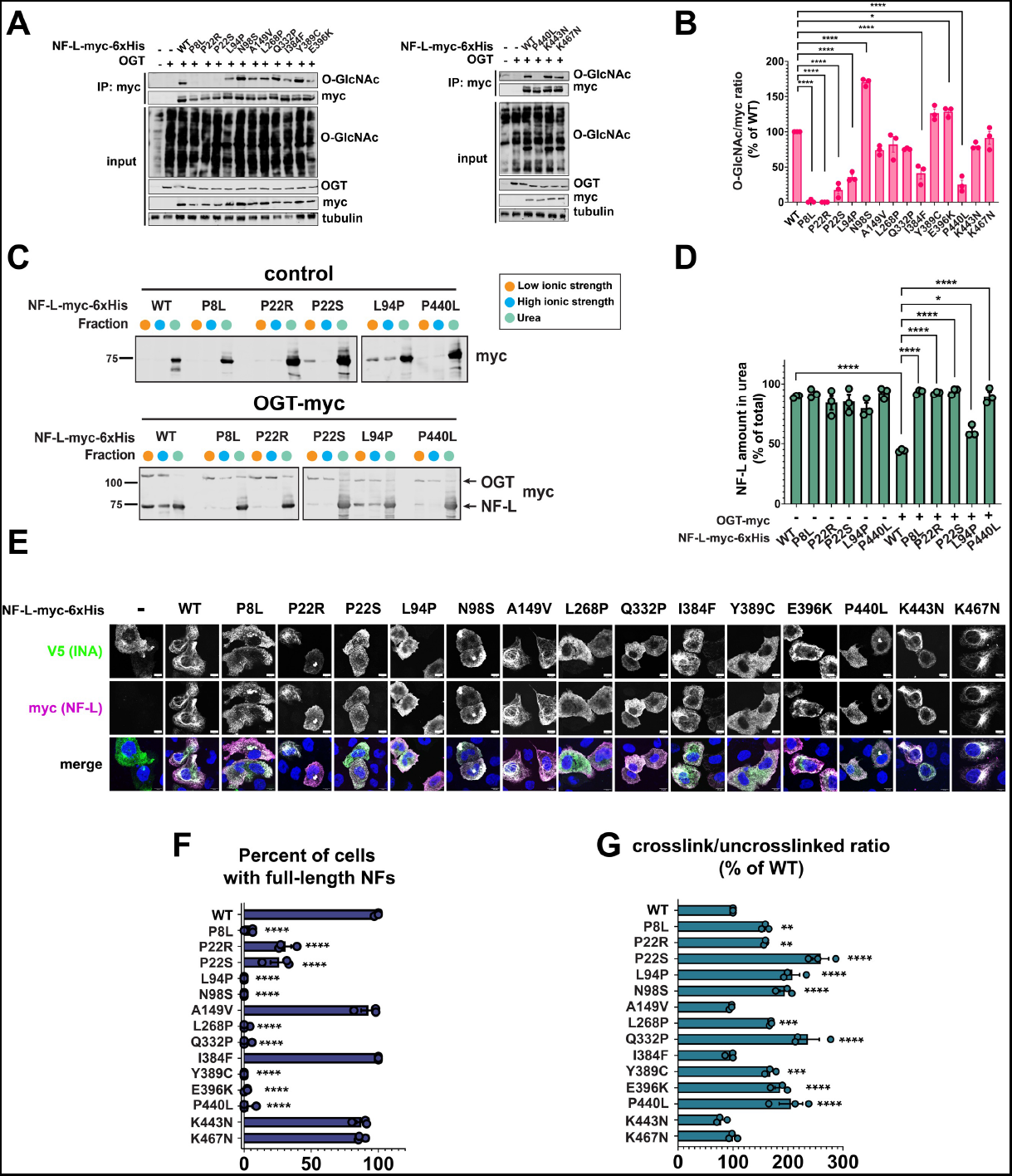
NF-L O-GlcNAcylation is dysregulated by CMT-causative mutations. (A) *NEFL*^-/-^ 293T cells were transfected with WT or CMT mutant NF-L-myc-6xHis + OGT for 24 hrs and analyzed by myc IP and IB. (B) Normalized O-GlcNAc signal (O-GlcNAc/myc ratio) was calculated for the experiments performed in (A) (n=3). (C) *NEFL*^-/-^ 293T cells were transfected with WT or CMT mutant NF-L-myc-6xHis ± OGT for 24 hrs and analyzed by differential extraction and IB. (D) NF-L amount extracted into urea buffer was calculated as percent of total NF-L across three fractions from the experiment described in (C) (n=3). (E) SW13 vim^-^ cells were transfected with WT or CMT mutant NF-L-myc-6xHis + INA-V5 for 24 hrs and analyzed by IFA. Scale bar: 10 µm. (F) Quantification of percent of cells with full-length NFs from experiment described in (E) was performed by a blinded researcher (n=3). (G) *NEFL*^-/-^ 293T cells were transfected with WT or CMT mutant NF-L-myc-6xHis ± 100 µM GlcNDAz for 48 hrs, subjected to UV crosslinking, and analyzed by IB. Normalized crosslink signal (crosslink/uncrosslinked ratio) was calculated (n=3). Throughout: *, p < 0.05; **, p < 0.005; ***; p < 0.0005; ****, p < 0.0001.

**Figure 7:**
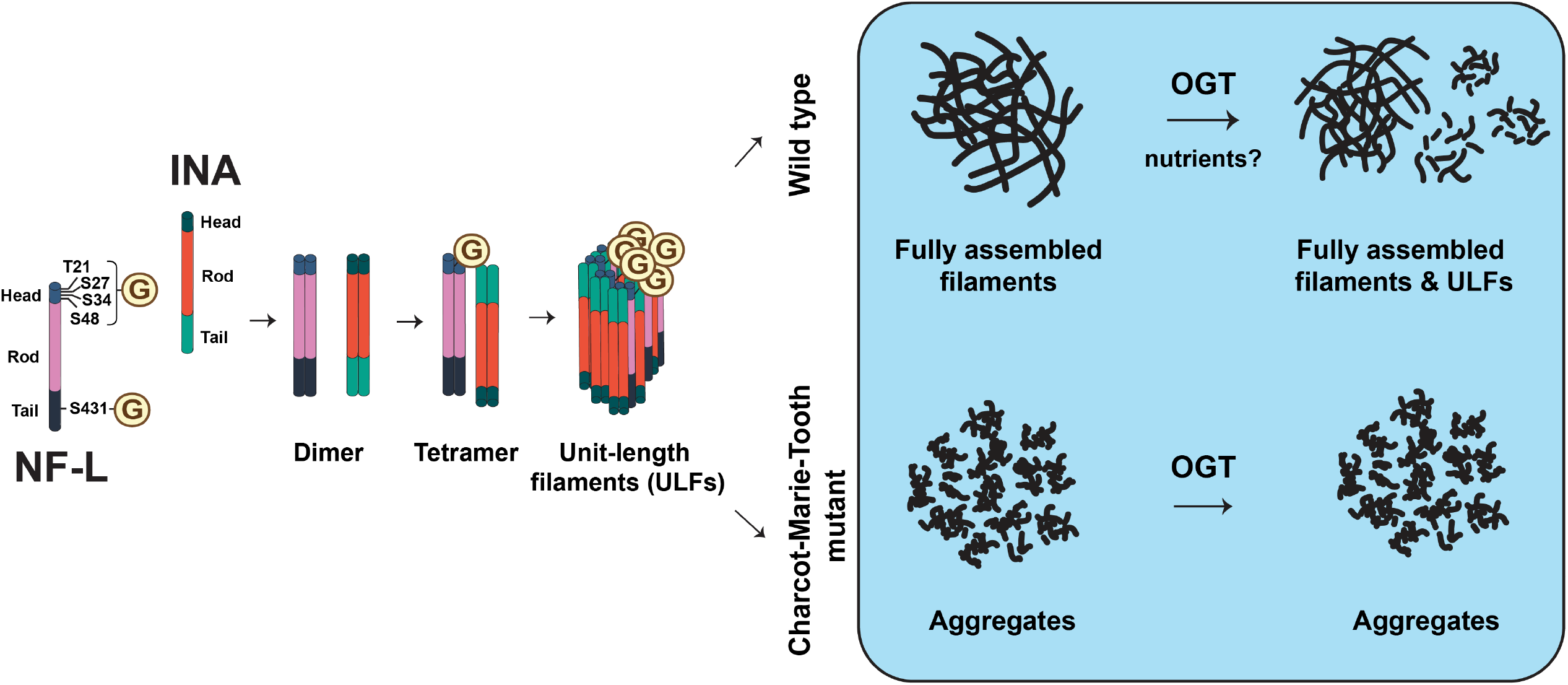
Site-specific O-GlcNAcylation regulates NF-L assembly and function and is perturbed by CMT disease mutations. Based on our results, we propose a model wherein nutrient-responsive O-GlcNAcylation of the NF-L head domain (T21/S27/S34/S48) promotes homotopic NF-L/NF-L and heterotypic NF-L/INA interactions and assembly and/or maintenance of assembled filaments under homeostatic conditions. Elevating NF-L O-GlcNAcylation reduces the prevalence of full-length filaments of WT NF-L, whereas CMT-causative mutations proximal to NF-L glycosites display lower O-GlcNAcylation and remain insensitive to the influence of OGT. The stoichiometry and topology of O-GlcNAc-mediated NF-L/INA interactions remain uncertain. One plausible model is depicted for simplicity.

## Discussion

Nervous system function depends on the unique sizes, morphologies, and intricate subcellular organization of individual neurons^1,5-8^. These properties, in turn, rely on a complex and dynamic neuronal cytoskeleton^1,5-8^. In particular, the NF network is required for neuronal shape, mechanical integrity, organelle trafficking, and synapse architecture^2-4^. In addition to their physiological importance, the dysregulation of NF proteins contributes to myriad neurological diseases^21,22,25,31-34^. Therefore, understanding the dynamic regulation of the NF cytoskeleton is a crucial – but incompletely realized – goal.

Here we show that NF-L, which is essential for NF formation^2-4^, is regulated by site-specific O-GlcNAcylation in cultured cells, primary neurons, and human brain tissue. Our MS site-mapping and mutagenesis data pinpointed five O-GlcNAc sites on human NF-L, predominantly in the head domain (Figure 1F). These results agree with prior proteomics studies of rodent NF-L orthologs^47-51^ and post-mortem human brain samples^52^, and we have extended this earlier work by validating and characterizing the function of human NF-L O-GlcNAcylation.

The regulation of human NF-L by O-GlcNAc is likely even more extensive than our current results indicate. For example, consistent with prior work^52^, our MS data detected an additional high-confidence O-GlcNAcylated peptide in the NF-L tail domain (residues 404-416), but the specific glycosite was not identifiable, due to low abundance. (Complete MS site-mapping data are provided – please see Materials and Methods for details.) Therefore, NF-L O-GlcNAcylation probably occurs on more residues than we have characterized thus far, perhaps explaining the residual O-GlcNAc signal detected on the NF-L^5A^ mutant (Figure 1H and J). In addition, our results on human NF-L may be relevant to other NF proteins and to other species. For example, some NF-L glycosites characterized here are conserved among vertebrate NF-L orthologs, and cognate residues in NF-medium (NF-M) (e.g., T19 for NF-L T21 and T48 for NF-L S48) are reportedly O-GlcNAcylated in mouse neurons^47,48^. Beyond NFs, we^81^ and others^96,97^ have previously demonstrated the functional importance of O-GlcNAcylation on the assembly states and functions of other IF proteins, implying that O-GlcNAc may be a general regulator of the > 70-member human IF protein family. Given this, thorough characterization of O-GlcNAcylation on NF-L and other NF proteins may provide insight into IF protein regulation in other tissues.

Our results also align with prior studies demonstrating the regulation of NF proteins by PTMs in general^41-46^. For example, NF proteins are among the most highly phosphorylated substrates in the brain^2^. Phosphorylation of the NF-L head domain regulates NF assembly^42,43^ and NF transport towards dendrites^44^, and phosphorylation of the tail domains of NF-M and NF-heavy (NF-H) impacts the regular spacing between assembled NFs^45,46^. Aberrant NF phosphorylation is a pathological feature of several human neurodegenerative diseases, including CMT type 2E^98^, AD^99^, and ALS^100^. Our results show that O-GlcNAcylation, like phosphorylation, can influence NF-L assembly state and downstream functions, such as the regulation of organelle transport (Figures 3). Interestingly, there is complex and well-documented crosstalk between phosphorylation and O-GlcNAcylation in many cellular contexts^101-106^. For example, these PTMs can compete for identical or nearby residues as modification sites, or, alternatively, can promote the modification of one residue by one PTM via the modification of a nearby residue by the other^101-106^. Additionally, OGT, OGA, kinases, and phosphatases modify each other and are often detected together in multiprotein complexes, constituting another layer of crosstalk^101,106^. However, the potential interplay between phosphorylation and O-GlcNAcylation on NF-L remains unknown. None of the NF-L glycosites that we identified is a known phosphorylation site^107-110^ and IP/IB experiments did not show detectable changes in NF-L phosphorylation when O-GlcNAcylation was manipulated or vice versa (data not shown). Examining potential phosphorylation-glycosylation crosstalk on NF proteins in other experimental contexts will be an important goal of later studies.

Functionally, we demonstrate that NF-L O-GlcNAcylation is required for the regulation of mitochondrial and lysosomal motility (Figure 3). Our results are consistent with prior reports of NFs influencing mitochondrial trafficking^16,24,26,28^. For example, in human iPSC-derived motor neurons, mitochondrial movement is increased in the absence of NF-L^16^ and decreased in the presence of NF-L mutants that aggregate in the soma^26^. In our experiments, the expression of WT NF-L suppressed mitochondrial motility, but expression of the NF-L^4A^ glycosite mutant, which exhibits assembly state defects (Figure 2E-F), failed to do so (Figure 3A-B). The endogenous microtubule cytoskeleton is intact under these conditions (Figure s3), indicating that gross microtubule disruption cannot explain the mutant phenotype. Instead, we propose that WT NF-L incorporates into the endogenous NFs, creating a robust network in the axonal cytoplasm that mitochondria must circumnavigate, whereas loss of NF-L O-GlcNAcylation dysregulates NF assembly state, reducing these barriers and accelerating mitochondria motility. Perhaps counter-intuitively, WT NF-L expression accelerated lysosomal motility in our system, whereas NF-L^4A^ slowed it (Figure 3C-D). It may be that comparatively small lysosomes benefit from less crowding by impeded mitochondria in the presence of a more elaborate WT NF network and yet are slowed via a distinct mechanism when O-GlcNAcylation of NF-L is reduced, but this hypothesis remains to be tested. Though one prior report found no effect of NF-L loss on lysosomal motility^16^, differences in experimental systems may account for this ostensible discrepancy with our work. Regardless, our data consistently indicate that NF-L O-GlcNAcylation impacts organelle motility in primary neurons. Mitochondria and lysosomes are transported along microtubules in the forward direction by kinesins and in the reverse by dyneins^88^. In principle, NF-L glycosylation could affect the action of one or both motor complexes. Consistent with our proposed model of passive hindrance of organelle motility by the WT NF network, our preliminary data suggest that anterograde and retrograde organelle motion are equally affected by the loss of NF-L O-GlcNAcylation (data not shown). However, further work will be required to determine whether kinesin-and dynein-dependent motility are differentially affected by NF O-GlcNAcylation in other contexts or model systems.

Mechanistically, earlier studies proposed a role for O-GlcNAcylation in regulating rodent NF assembly^47,48^. Our results support this hypothesis by demonstrating that O-GlcNAc impacts human NF-L function at least in part by modulating its assembly state and PPIs. Biochemical and imaging experiments showed that O-GlcNAcylation drives WT NF-L to lower-order assembly states, reducing the prevalence of full-length NFs (Figure 2). Furthermore, our GlcNDAz crosslinking experiments showed that NF-L engages in both homotypic (NF-L/NF-L) and heterotypic (NF-L/INA) O-GlcNAc-mediated interactions (Figure 5), providing a potential mechanism to explain the effects of NF-L O-GlcNAcylation on NF assembly state (Figure 7). Taken together, these results revealed previously unknown, glycan-mediated biochemical interactions among NF proteins and provide important new information on the molecular regulation of the NF cytoskeleton.

Despite these insights, additional questions remain. For example, the stoichiometry and structural basis of O-GlcNAc-mediated NF-L PPIs are as yet unknown. Based on the apparent molecular weight of GlcNDAz-dependent crosslinks (∼250 kDa) and the evidence for homotypic NF-L/NF-L PPIs (Figure 5D), we propose that these complexes contain four or five distinct polypeptides, including at least two NF-L (62 kDa) and at least one INA (55 kDa) molecules. Because increased O-GlcNAcylation potentiates NF-L/INA interactions (Figure 5E-G) and reduces the prevalence of fully assembled NFs (Figure 2B-F), we hypothesize that O-GlcNAc-mediated homotypic and heterotypic PPIs of NF-L occur within lower-order assembly states *in vivo*. We demonstrated that ablating all five identified human NF-L glycosites significantly decreased NF-L/INA interactions (Figure 5F-G), indicating that O-GlcNAcylation of NF-L itself is at least partly responsible for these interactions. Whether INA is also O-GlcNAc-modified has not been examined specifically, but untargeted glycoproteomics studies have reported data suggesting that it might be^50-52,111,112^. O-GlcNAc moieties on INA could hypothetically contribute to interactions with NF-L. Because INA exhibits aberrant PPIs in NF inclusion body disease^93,94^, these questions may have clinical relevance. However, the paucity of information on INA O-GlcNAcylation and the total lack of structural information on NF proteins^113^ currently hampers a more detailed characterization of these interactions. Additional biophysical and MS experiments in future studies will be needed to define the stoichiometry and structural underpinnings of O-GlcNAc-mediated PPIs among NF components.

Our results indicate that upstream signals could regulate NF-L by inducing or inhibiting its O-GlcNAcylation (Figures 1-3, s1-2), and we identify fluctuations in nutrient or growth factor availability as candidate stimuli that may govern NF-L O-GlcNAcylation *in vivo* (Figure 4). Although further studies will be needed to dissect the relationship between nutrient-sensing and NF-L glycosylation in neurons, similar phenomena have been reported in other systems. For example, the Hart lab previously showed that glucose deprivation alters the O-GlcNAcylation of NF-H in a p38 mitogen-activated protein kinase-dependent manner, leading to increased NF-H solubility^114^. In other contexts, the Schwarz lab demonstrated that local glucose concentration differences within the axon regulate mitochondrial motility through the O-GlcNAcylation of microtubule-dependent motor complex components^115^. Experiments are currently underway to determine the impact of nutrient and growth factor changes on NF-L O-GlcNAcylation and function, including organelle motility, in cultured primary neurons or human iPSC-derived neurons.

Finally, our work may have implications for neurodegenerative disorders characterized by NF protein aggregation, particularly CMT. Previous studies of CMT-causative NF-L mutants demonstrated their assembly defects in cultured cells^29^, mouse brains^27^, and iPSC-derived motor neurons^28^. In agreement with these results, most of the CMT NF-L mutants we tested abrogated filament formation (Figure 6E-F). Strikingly, we also found that CMT mutations near glycosites (e.g., P8L, P22R, P22S, P440L) significantly reduced or entirely ablated NF-L O-GlcNAcylation (Figure 6A-B) and abolished the OGT-induced shift to lower-order assembly states exhibited by WT NF-L (Figure 6C-D). These CMT mutations may alter the primary sequence or secondary structural determinants required by OGT to bind NF-L and/or may adopt abnormal tertiary or quaternary structures that occlude OGT’s access to its target residues. Consistent with this notion, a recent study showed that the NF-L head domain, which is rich in low-complexity sequence^116^, self-associates via labile cross-β structures^116^. Residues P8 and P22, which are CMT mutational hotspots near glycosites^21^, reduce the formation of cross-β structures^116^, and mutations at these positions result in enhanced polymerization and head-domain self-association^116^. It will be interesting to determine in future work whether loss of O-GlcNAcylation on these CMT mutants promotes their adoption of aberrant conformations, perhaps leading to aggregation and downstream pathological effects.

A longstanding puzzle in the CMT field is that mutations all along the NF-L protein can cause the disease, yet no clear correlations have emerged between *NEFL* genotypes and molecular, cellular, or clinical phenotypes^21,22^. In some respects, our results reflect a similar conundrum. For instance, while the abovementioned mutations near glycosites reduce NF-L O-GlcNAcylation, other CMT mutants showed elevated glycosylation, relative to WT (Figure 6A-B). Similarly, while most CMT mutants show greatly reduced filament formation, others (e.g., A149V, I384F, K443N, K467N) appear indistinguishable from WT in this regard (Figure 6E-F). Finally, most CMT mutants demonstrated enhanced levels of GlcNDAz crosslinking, relative to WT (Figure 6G), even though several of these also show reduced levels of O-GlcNAcylation (Figure 6A-B). It may be that particularly aggregation-prone CMT mutants adopt irregular conformations^21^ that predispose them to enhanced, non-specific crosslinking in the GlcNDAz assay. In support of this model, we showed that the CMT mutants that can still form filaments (A149V, I384F, K443N, K467N) also exhibit near-WT levels of GlcNDAz crosslinking (cf. Figure 6E-F and 6G).

Additional studies will be needed to dissect the mechanistic explanations for these observations. Nevertheless, our results consistently indicate that most CMT mutations cause anomalies in NF-L O-GlcNAcylation, assembly state, and PPIs. In future work, it will be important to determine whether dysregulated O-GlcNAcylation of NF-L or other NF proteins contributes to their aggregation and to the subsequent pathological effects in CMT and other neurodegenerative disorders, analogous to the disease-associated hyperphosphorylation of NF-M and NF-H^98-100^. Recent pre-clinical studies and clinical trials have demonstrated that newly developed small molecule inhibitors can cross the blood-brain barrier, potently modulate global O-GlcNAc levels, and ameliorate some aspects of neurological diseases^62,63^. Sensitive assays of NF-L in CSF have also shown great promise as biomarkers for a range of nervous system conditions^35^. In this context, our results may point to future opportunities to exploit particular glycoforms of NF-L as diagnostic biomarkers or to rationally manipulate NF protein O-GlcNAcylation to correct pathological aggregations in CMT and other neurological disorders.

## Acknowledgements

We thank Dr. So Young Kim (Duke) for assistance with genome engineering and generation of the pSpCas9-GFP-*AAVS1* gRNA construct, Dr. Bin Li (Duke) for help with single-cell sorting, Dr. Chao-Chieh Lin (Duke) for help with qPCR, Noah Linhart (Duke) for help with GlcNDAz proteomic sample preparation, and Dr. Natasha Snider (UNC Chapel Hill) for SW13 vim^-^ cells. This work was supported by US National Institutes of Health (NIH) grant R01GM118847 to M.B., NIH grant R01NS111588 and a gift from the Hannah’s Hope Foundation to M.B. and J.T.C., a Research Award for Graduate Students from the Ruth K. Broad Biomedical Research Foundation to D.T.H., a Hannah Gray Fellowship from the Howard Hughes Medical Institute and support from the Duke Science and Technology Scholars program to C.S.E., and NIH Alzheimer’s Disease Research Center grant P30AG072958 to the Duke Bryan Brain Bank and Biorepository.

## Contributions

M.B. and D.T.H. conceived and supervised the project. D.T.H., J.H., K.N.T., and A.J.W. performed the experiments. J.R.S. and C.S.E. prepared the rat hippocampal neurons and consulted on organelle motility assays and data analysis. E.J.S. performed the MS-identification of human NF-L O-GlcNAc sites and GlcNDAz proteomics. J.T.C. consulted on the strategic directions of all experiments. M.B. and D.T.H. wrote and all authors reviewed the manuscript.

## Figure legends

**Figure s1:**
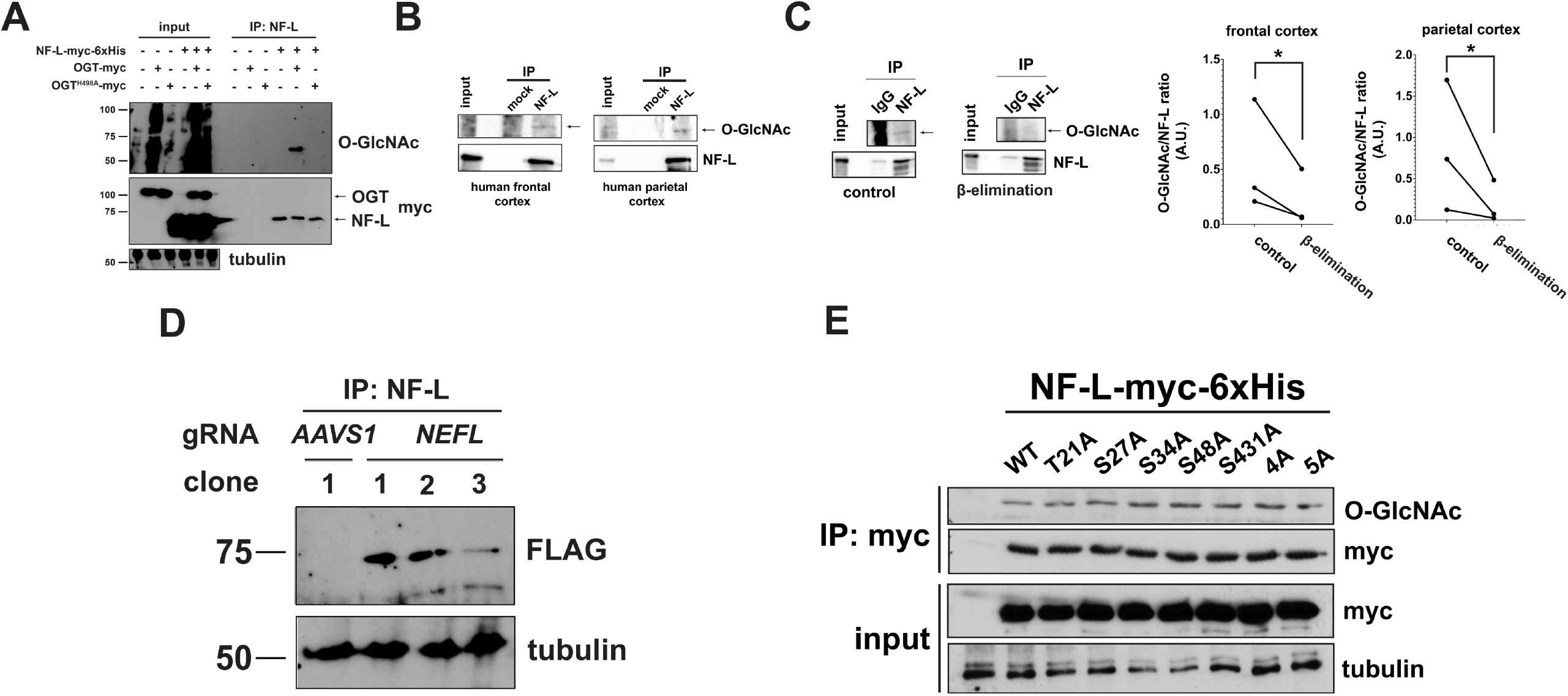
Site-specific O-GlcNAcylation of the human NF-L head and tail domains. (A) 293T cells were transfected with NF-L-myc-6xHis ± OGT-myc or OGT^H498A^-myc for 24 hrs, and lysates were analyzed by NF-L IP and IB. (B) Human frontal or parietal cortex homogenates were analyzed by NF-L IP and IB. (C) Human frontal or parietal cortex homogenates were analyzed by NF-L IP, on-blot β-elimination, and IB. Left: Representative frontal cortex IP/IBs. Right: Normalized O-GlcNAc signal (O-GlcNAc/NF-L ratio) was calculated (n=3). *, p < 0.05. (D) Lysates from single cell-derived clones of endogenously tagged NF-L-3xFLAG-6xHis 293T cells or negative control were analyzed by NF-L IP and IB. (E) *NEFL*^-/-^ 293T cells were transfected with WT or glycosite mutant NF-L-myc-6xHis for 24 hrs, and lysates were analyzed by myc IP and IB.

**Figure s2:**
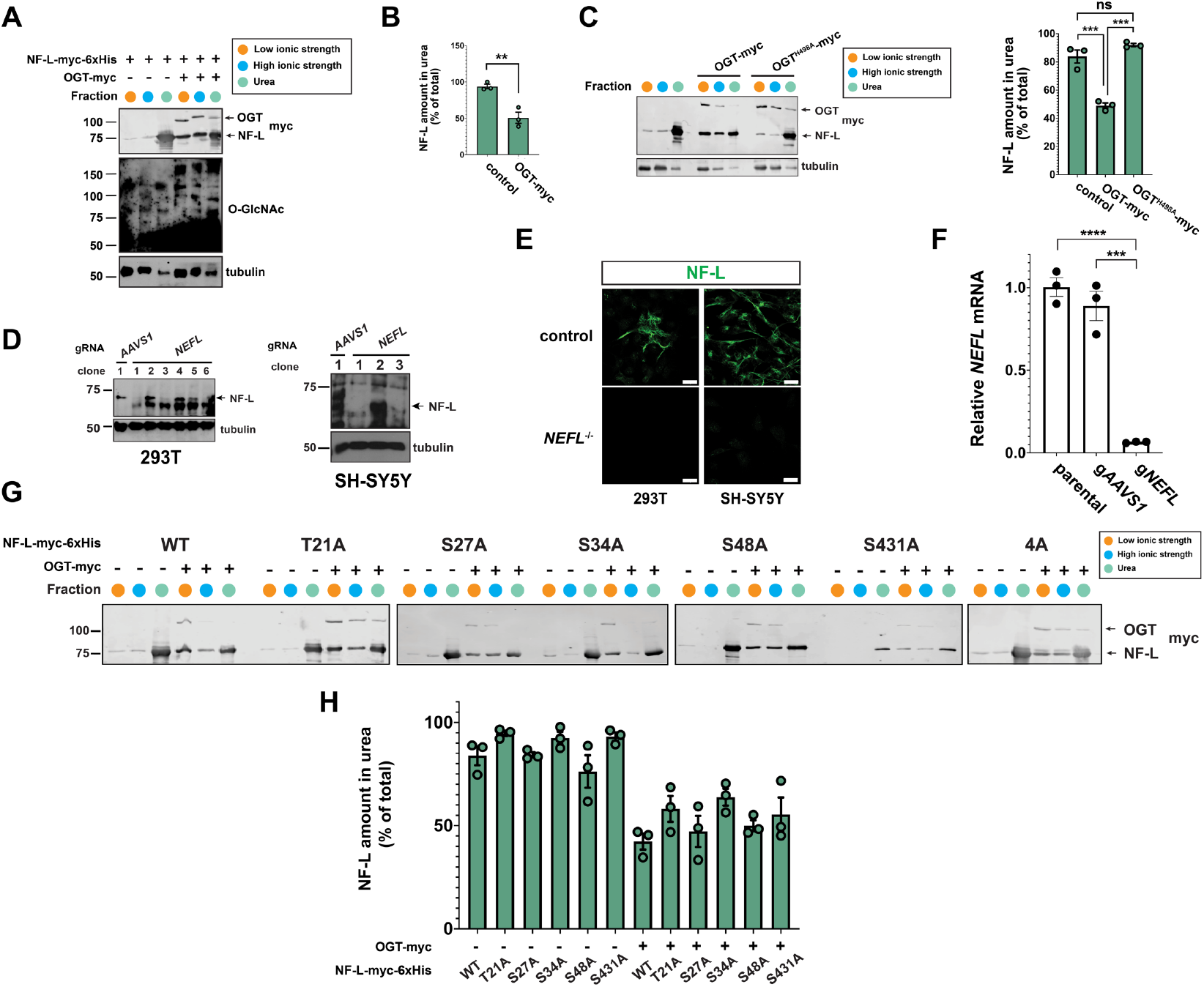
NF-L O-GlcNAcylation influences NF-L assembly state and filament formation. (A) SH-SY5Y cells were transfected with NF-L-myc-6xHis ± OGT-myc for 24 hrs and analyzed by differential extraction and IB. (B) NF-L amount extracted into urea buffer was calculated as percent of total NF-L across three fractions from the experiment described in (A) (n=3). (C) Left: 293T cells were transfected with NF-L-myc-6xHis ± OGT-myc or OGT^H498A^-myc for 24 hrs and analyzed by differential extraction and IB. Right: NF-L amount extracted into urea buffer was calculated as percent of total NF-L across three fractions (n=3). (D-F) Successful CRISPR/Cas9-mediated deletion of *NEFL* in a single cell-derived clone was verified by IB (D), IFA (Scale bar: 20 µm) (E), and actin-normalized qPCR (F). An *AAVS1* gRNA serves as a negative control for CRISPR/Cas9 deletion. mRNA from parental SH-SY5Y cells serves as a positive control for qPCR (F). (G) *NEFL*^-/-^ 293T cells were transfected with WT or glycosite mutant NF-L-myc-6xHis ± OGT-myc for 24 hrs and analyzed by differential extraction and IB. (H) NF-L amount extracted into urea buffer was calculated as percent of total NF-L across three fractions from the experiment described in (G) (n=3).

**Figure s3:**
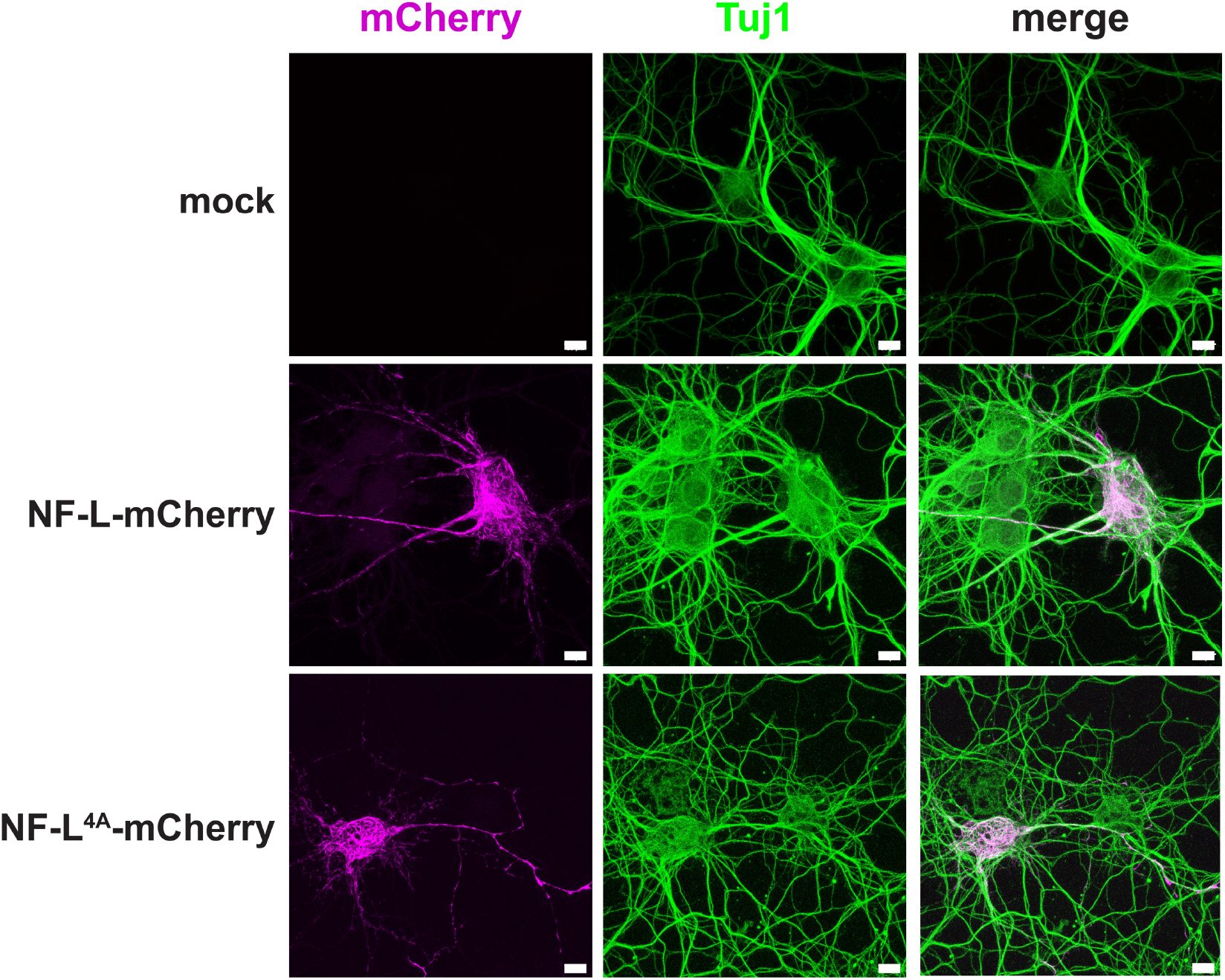
NF-L expression does not disrupt the neuronal microtubule cytoskeleton. Cultured E18 rat hippocampal neurons at day 6 *in vitro* were transfected with WT or NF-L^4A^-mCherry and analyzed by IFA with Tuj1 and mCherry. Scale bar: 10 µm.

**Figure s4:**
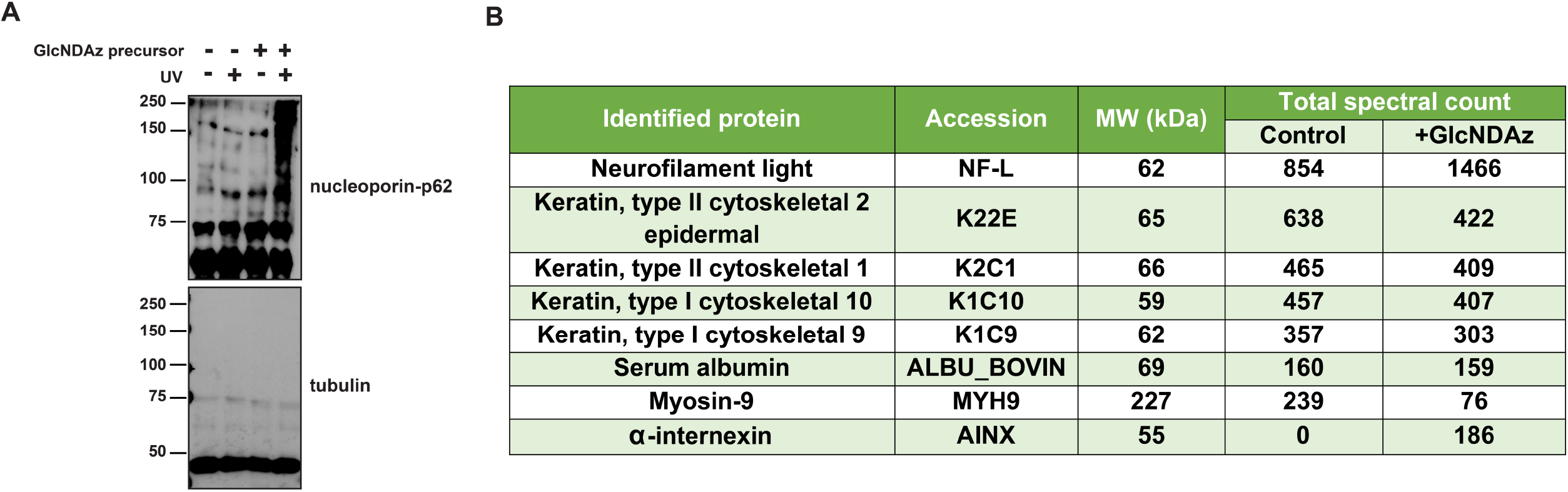
Direct, O-GlcNAc-mediated interactions between NF-L and INA. (A) 293T cells were treated with DMSO (vehicle) or 100 µM GlcNDAz for 48 hrs, subjected to UV crosslinking, and analyzed by IB. Heavily glycosylated nucleoporin-62 is a positive control, whereas unglycosylated tubulin is a negative control. (B) 293T cells were transfected with NF-L-myc-6xHis ± 100 µM GlcNDAz for 48 hrs and subjected to UV crosslinking. Lysates were analyzed by tandem myc IP/Ni-NTA purification, SDS-PAGE, and colloidal blue staining. High molecular weight crosslinked NF-L complexes from ± GlcNDAz samples were excised from the gels and analyzed by MS proteomics. Top protein IDs are shown. INA was identified from GlcNDAz-treated cells, whereas no INA peptides were detected in the corresponding gel region from DMSO (vehicle)-treated cell samples. All IDs except NF-L and INA are common contaminants.

## Materials and Methods

### Chemicals

Thiamet-G was purchased from Cayman Chemical (#13237). Peracetylated 5SGlcNAc and Ac_3_GlcNDAz-1P(Ac-SATE)_2_ were synthesized by the Duke Small Molecule Synthesis Facility essentially as described^77,91^.

### DNA constructs

The mito-mEmerald and LAMP1-GFP constructs have been described^85^. NF-L-V5/plenti6.3-DEST was purchased (DNASU, HsCD00870013). Primers were designed with QuikChange Primer Design (Agilent). To make NF-L-myc-6xHis glycosite mutants, NF-L-mCherry, OGT-3xFLAG, INA-V5, and UAP1^F383G^-FLAG constructs, PCR was performed using Phusion Hot Start II DNA polymerase (Thermo Fisher Scientific, F549S) with primers included in Table 1. PCRs were digested with DpnI (New England Biolabs [NEB], R0176S), purified using a gel DNA recovery kit (Zymo Research, D4002), and ligated using NEBuilder HiFi DNA Assembly Cloning Kit (NEB, E5520S) with 1:3 vector:insert mass ratio calculated by NEBioCalculator (https://nebiocalculator.neb.com/#!/ligation). Then, the ligated product was transformed in 10-beta competent *E. coli* (NEB, C3019H), and colonies were picked for maxipreps (ZymoPURE II Plasmid Purification Kit, Zymo Research, D4202) and Sanger sequencing. Guide RNAs (gRNAs) for *NEFL* knockout (5’ – CTCGTAGCTGAAGGAACTCA – 3’) and *NEFL* knock-in (5’ – GTAGCTGAAGGAACTCATGG – 3’) were designed by DESKGEN tool and subcloned into pSpCas9(BB)-2A-GFP (Addgene, 48138) with BbsI-HF (NEB, R3539) and T4 DNA ligase (NEB, M0202S). For the *NEFL* knock-in repair template, the 3xFLAG-6xHis gene fragment (IDT) was subcloned into the *AAVS1*_3xFLAG-2xStrep plasmid (Addgene, 68375) at the NcoI/BstBI sites (“*AAVS1*_3xFLAG-6xHis”). The *NEFL* homology arm (HA)_1 fragment (Genscript) was cloned into the *AAVS1*_3xFLAG-6xHis vector at the Nde1/NcoI sites. The HA_2 fragment (Genscript) was then ligated to the *AAVS1*_3xFLAG-6xHis-HA_1 vector at the BstBI/EcoRI sites. The nucleotide sequences for the NF-L^4A^ gene fragment (IDT), 3xFLAG-6xHis, and two HA fragments are included in the Table 2.

**Table 1:**
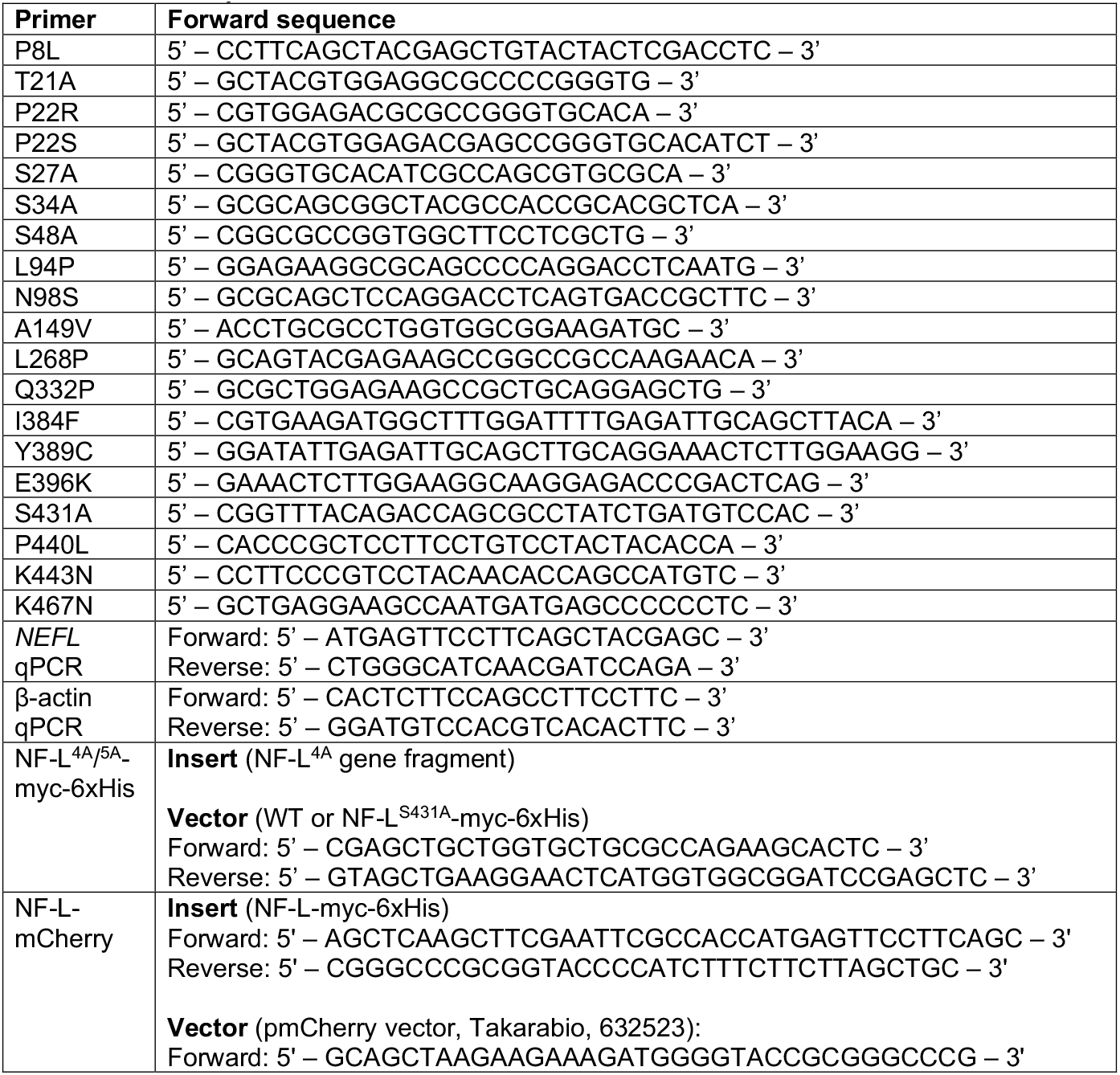
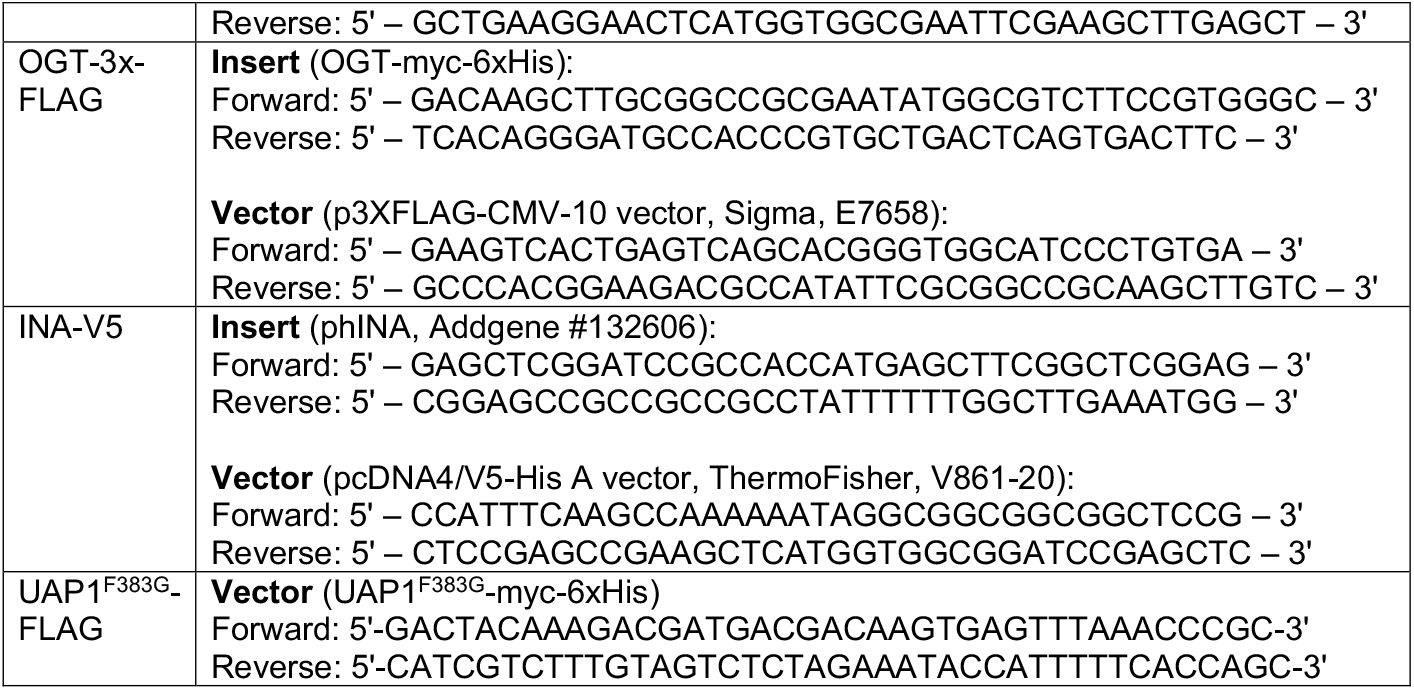
Primer sequences

**Table 2:**
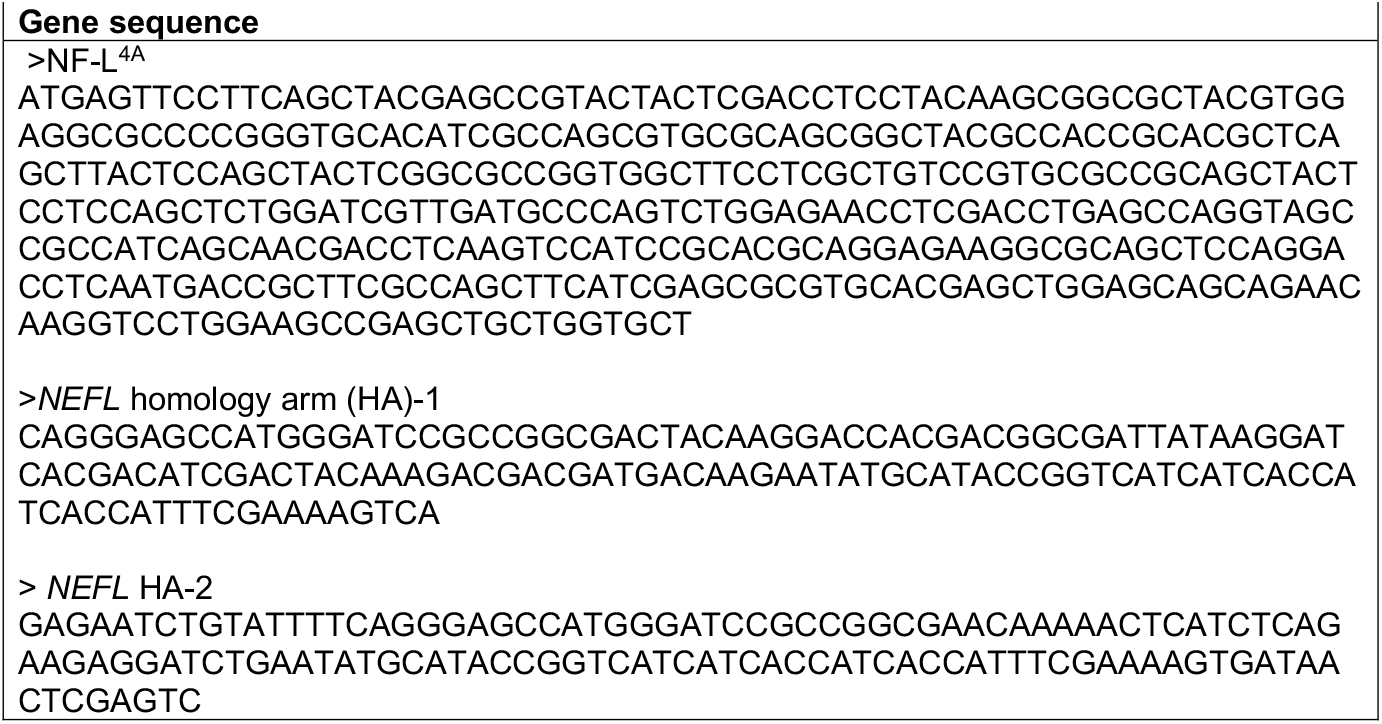
Gene blocks and fragments

### Cell culture

293T (ATCC CRL-11268) and SW13 vim^-^ cells (Snider lab, UNC) were grown in Dulbecco’s modified Eagle Medium (DMEM, Sigma Aldrich, D6429) containing 10% fetal bovine serum (FBS, Sigma-Aldrich, F0926), 100 units/mL penicillin, and 100 µg/mL streptomycin (Pen/Strep, Gibco, 15140-122). SH-SY5Y (ATCC CRL-2266) cells were grown in DMEM/F12 (Gibco, 11330-032) containing 10% FBS and 1% Pen/Strep. Following flow cytometry for CRISPR manipulations, sorted, single GFP-positive cells were plated into DMEM (293T) or DMEM/F12 (SH-SY5Y) with 15% FBS, 1% Pen/Strep. For sorted SH-SY5Y cells, DMEM/F12 was changed every two days. For nutrient starvation, *NEFL*^-/-^ 293T cells expressing NF-L-myc-6xHis were starved for 48 hrs of glucose: glucose-free DMEM (Thermo Fisher Scientific, 11966025) with 1 mM sodium pyruvate (Thermo Fisher Scientific, 11360070), 10% FBS, and Pen/Strep; glutamine: glutamine-free DMEM (Thermo Fisher Scientific, 11960044) with 1 mM sodium pyruvate, 10% FBS, and Pen/Strep; or serum: DMEM with 1 mM pyruvate and Pen/Strep. For hippocampal neurons, embryonic day 18 Sprague Dawley rat hippocampi were isolated as described^85^. Neurons were plated on 35 mm culture dishes precoated with 0.5 mg/mL poly-L-lysine (Sigma Aldrich, P1274) at a density of 125,000 cells/dish. Neurons were initially plated in minimum essential medium (Gibco, 11095-072) supplemented with 10% horse serum (Gibco, 16050122), 33 mM D-glucose (Sigma, G8769-100), and 1 mM sodium pyruvate (Corning, 25-000Cl) and incubated for 2-5 hrs. The medium was then replaced with Neurobasal (Thermo Scientific, 21103049) supplemented with 33 mM D-glucose, 2 mM GlutaMAX (Life Technologies, 35050061), 1% Pen/Step, and 2% B-27 (ThermoFisher, 12-587-010). Cytosine arabinoside (5 µM) was added the day after plating to prevent glial cell proliferation. All cell lines were maintained at 37 °C in a 5% CO_2_ atmosphere.

### Transfections

Unless otherwise indicated, cells were transfected with 10 µg of DNA at 70-80% cell density using TransIT-LT1 (Mirus, 2300) for 293T cells and at 50-60% cell density with Lipofectamine 3000 (ThermoFisher Scientific, L3000001) for SH-SY5Y cells. For co-IPs, 293T cells were transfected with 8 µg of each DNA at 25-30% cell density. For CRISPR-tagging of *NEFL*, 10 million 293T cells seeded on a 15 cm culture dish were transfected with 10 µg of pSpCas9-GFP-*NEFL* knock-in construct (or pSpCas9-GFP-*AAVS1*, negative control) single gRNA (gRNA) and 20 µg of *NEFL* homology-directed repair vector at 50-60% cell density. For CRISPR deletion, 10 million 293T or 10 million SH-SY5Y cells seeded in a 15 cm culture dish were transfected with 15 µg of pSpCas9-GFP-*NEFL* knockout construct (or pSpCas9-GFP-*AAVS1*, negative control) gRNA.

### IP/co-IP

24 hrs (or 48 hrs for co-IPs) post-transfection, cells were harvested in cold phosphate-buffered saline (PBS) and lysed in cold IP lysis buffer (20 mM Tris-HCl pH 7.4, 1% Triton X-100, 0.1% SDS, 150 mM NaCl, 1 mM EDTA) with protease inhibitor cocktail (Sigma, P8340, 1:100), 200 µM Na_3_VO_4_ (Millipore Sigma, 13721-39-6), 50 µM UDP (Sigma, 94330; OGT inhibitor), and 5 µM PUGNAc (Cayman Chemical, 17151; OGA inhibitor). Lysates were incubated on ice for 15 min, probe-sonicated for 50 s at 40% duty cycle and cleared by centrifugation at 27,000 g for 15 min at 4 °C. Cleared lysates were quantified by bicinchoninic acid (BCA) assay (ThermoFisher, 23225). IPs/co-IPs were performed on 1.5-2 mg of total protein in 0.5 mL (∼3-4 mg/mL protein). Unless otherwise indicated, 3 µg of primary antibody per 1 mg of protein was added to the protein lysate for rotation overnight at 4 °C. The next day, 20 µL of settled protein A/G UltraLink resin (ThermoFisher, 53133) were washed three times in the corresponding lysis buffer and added to each IP/co-IP for rotation at room temperature (RT) for 2 hr. Proteins were eluted in lysis buffer supplemented with 5% fresh β-mercaptoethanol (Sigma, M3148) and 1X SDS-PAGE loading buffer (5X SDS-PAGE loading buffer: 250 mM Tris pH 6.8, 10% SDS, 30% glycerol, 5% β-mercaptoethanol, 0.02% bromophenol blue). Eluates were heated at 95 °C for 5 min and analyzed by IB.

### IB

For enhanced chemiluminescence (ECL) detection, SDS-PAGE gels were electroblotted onto 100% methanol pre-soaked polyvinylidene difluoride membranes (PVDF, 0.45 µm, Thermo Fisher Scientific, 88518) in transfer buffer (25 mM Tris pH 8, 192 mM glycine, 0.1% SDS, 20% methanol) using a BioRad TransBlot Turbo system. Then, membranes were incubated in blocking buffer (2.5% (w/v) bovine serum albumin [BSA] in Tris-buffered saline with Tween [TBST] [20 mM Tris-HCl pH 7.4, 150 mM NaCl, 0.1% Tween 20]) with agitation at RT for 30 min. Membranes were incubated with primary antibody diluted 1:1,000 (except for α-tubulin, 1:100,000) in blocking buffer overnight with gentle shaking at 4 °C. The next day, membranes were washed three times with TBST, each 10 min, and incubated with the appropriate horseradish peroxidase (HRP)-conjugated secondary antibody diluted 1:10,000 in blocking buffer at RT for 1 hr. Membranes were again washed three times with TBST, each 10 min. Bands were visualized using ECL (Genesee Scientific 20-300B) and photographic film (LabScientific, XAR ALF 2025). For quantitative fluorescent IBs, gels were electroblotted onto nitrocellulose membrane (0.45 µm, Bio-Rad, 1620115). Blocking, primary, and washing conditions were same as above. Membranes were incubated with appropriate IRDye-conjugated secondary antibody diluted 1:30,000 in blocking buffer at RT in the dark for 1 hr. Membranes were washed three times with TBS, each 10 min. Bands were visualized using a LI-COR Odyssey DLx Imaging System. Complete antibody information is provided in Table 3.

**Table 3:** Antibody inventory

| Antibody | Vendor and catalog number |
| --- | --- |
| myc (9E10) | Biologend, 626802 |
| O-GlcNAc (18B10.C7) | EMD Millipore, 05-1244 |
| O-GlcNAc (RL2) | Biologend, 677902 |
| O-GlcNAc (MultiMAb) | Cell Signaling Technology, 82332 |
| NF-L (C28E10) | Cell Signaling Technology, 2837 |
| FLAG (M-2) | Sigma Aldrich, F1804 |
| β-III tubulin (Tuj1) | R&D Systems, MAB1195-SP |
| α-tubulin | Sigma Aldrich, T6074 |
| V5 | ThermoFisher Scientific, R960-25 |
| V5 (D3H8Q) | Cell Signaling Technology, 13202 |
| INA | Novus, NB300-139 |
| Nucleoporin-p62 | BD Biosciences, 610498 |
| Goat HRP-conjugated anti-mouse IgG secondary antibody | SouthernBiotech, 1030-05 |
| Goat HRP-conjugated anti-mouse kappa secondary antibody | SouthernBiotech, 1050-05 |
| Goat HRP-conjugated anti-rabbit IgG secondary antibody | SouthernBiotech, 4030-05 |
| Goat HRP-conjugated anti-rabbit light-chain secondary antibody | SouthernBiotech, 4060-05 |
| Goat IRDye 800CW-conjugated anti-mouse IgG (H+L) secondary antibody | Li-Cor, 925-32210 |
| Goat IRDye 800CW-conjugated anti-rabbit IgG (H+L) secondary antibody | Li-Cor, 925-32211 |
| Goat Alexa Fluor 488-conjugated anti-mouse (H + L) secondary antibody | Thermo Fisher Scientific, A-11001 |
| Goat Alexa Fluor 488-conjugated–anti-rabbit (H + L) conjugated secondary antibody | Thermo Fisher Scientific, A-11008 |
| Goat Alexa Fluor 594-conjugated–anti-mouse (H + L) secondary antibody | Thermo Fisher Scientific, A-11005 |
| Goat Alexa Fluor 594-conjugated anti-rabbit (H + L) secondary antibody | Thermo Fisher Scientific, A-11012 |
| Goat Alexa Fluor 647-conjugated anti-mouse IgG (H+L) cross-adsorbed secondary antibody | Thermo Fisher Scientific, A-21235 |
| Goat Alexa Fluor 647-conjugated anti-rabbit IgG (H+L) Highly cross-adsorbed | Invitrogen, A-32733 |

### NF-L O-GlcNAcylation

For endogenous NF-L O-GlcNAcylation, SH-SY5Y cells cultured in 50 µM Thiamet-G were lysed in 8 M urea in PBS on ice for 30 min, homogenized by probe-sonication for 50 s at 40% duty cycle, and exchanged to IP lysis buffer using Zeba spin desalting columns (ThermoFisher Scientific, 89892). For brain NF-L O-GlcNAcylation, post-mortem human brain chips (Duke Bryan Brain Bank and Biorepository) were thawed on ice for 3-4 hrs, lysed in ice-cold 8 M urea in PBS by pipetting, and rotated for 20 min at 4 °C. Samples were probe-sonicated on ice with 40% duty cycle for 90 s, 50% for 90 s, and 60% for 30 s. Lysates were cleared by centrifugation at 27,000 g, 30 min at 4 °C, and the supernatant lipid layer was removed. Cleared lysates were transferred to a new microfuge tube. The remaining pellet was again extracted as above. The cleared samples were combined, exchanged into IP lysis buffer supplemented with protease inhibitor cocktail (1:100), 200 µM Na_3_VO_4_, 5 mM NaF, 50 µM UDP, and 5 µM PUGNAc using a Zeba column, and analyzed by IB.

### On-blot β-elimination

Brain homogenates were run in duplicate on an SDS-PAGE gel and electroblotted onto a single PVDF membrane as above. The membrane was cut in half vertically, with one half incubated in 55 mM NaOH and the other in water (control) at 40 °C with rocking, as described^78^. Membranes were then washed three times in TBS pH 7.4, blocked with 2.5% (w/v) BSA in TBST at RT for 1 hr, and analyzed by IB.

### CRISPR tagging or deletion of *NEFL*

48 hrs post-transfection with the constructs listed above, single GFP-positive cells were sorted into 96-well plates using a DiVa fluorescence-activated cell sorter (BD Biosciences) at the Duke Cancer Institute Flow Cytometry Shared Resource. Sorted cells were expanded into single cell-derived clones and verified for NF-L-3xFLAG-6xHis expression by reciprocal IP/IB or for *NEFL* deletion by quantitative RT-PCR, IP/IB, and IFA with NF-L antibody. Validated clones of NF-L-3xFLAG-6xHis-expressing 293T, *NEFL*^-/-^ 293T, and *NEFL*^-/-^ SH-SY5Y were selected for subsequent experiments.

### qPCR

Parental or *NEFL*^-/-^ SH-SY5Y cells seeded in a 6-well plate were collected at 80% cell density, lysed, and RNA-extracted using the RNeasy Mini Kit (Qiagen, 74104). RTL buffer was supplemented with β-mercaptoethanol prior to lysis. RNA concentration was measured by Nanodrop 2000 Spectrophotometer (ThermoFisher Scientific, ND-2000). *NEFL* cDNAs were made using SuperScript II reverse transcriptase II (Thermo Fisher Scientific, 18064014) and quantified by qPCR. The qPCR primer pairs (Table 1) were complementary to different regions of the *NEFL* mRNA. Relative levels of *NEFL* mRNA were normalized to β-actin mRNA in each sample. Triplicate reactions were performed using Sybr Master Mix (Life Tech-Power Sybr, 4367659) and a StepOnePlus Real-Time PCR system (Applied Biosystems).

### Endogenous NF-L purification for glycosite mapping

NF-L-3xFLAG-6xHis 293T cells were treated with 50 µM Thiamet-G and 4 mM glucosamine for 24 hrs as described^82,83^. Then, cells from 45 15 cm culture dishes were harvested, lysed, and quantified as for IP/co-IP. 270 mg total protein was divided evenly into 11 5-mL centrifuge tubes (Genesee Scientific, 24-285) and rotated with FLAG antibody overnight at 4 °C. 75 µL settled, washed protein A/G UltraLink resin was then added and rotated at RT for 2 hrs, then all tubes were pooled. Resin was washed with IP lysis buffer (20 mM Tris-HCl pH 7.4, 1% Triton X-100, 0.1% SDS, 150 mM NaCl) five times and rotated in 600 µL buffer A (300 mM NaCl, 1% Triton X-100, and 10 mM imidazole in 8 M urea/PBS) twice, each for 30 min, at 4 °C. After centrifugation (2,500 g, 5 min), the total cleared supernatant (1.2 mL) was rotated with 100 µL of HisPur Ni-NTA resin (ThermoFisher, 88223) for 4 hrs at 4 °C. Resin was then washed ten times with buffer A and eluted five times, each in 60 µL of elution buffer (250 mM imidazole in 8M urea/PBS) for 20 min with vigorous shaking. Purified NF-L samples were processed either from colloidal blue-stained (Thermo Fisher Scientific, LC6025) SDS-PAGE gel band or directly in-solution from the eluate.

### Liquid chromatography-tandem MS analysis

For gel band analysis, colloidal blue-stained SDS-PAGE bands (Invitrogen Unpaged 4-12% Bis-Tris) were manually excised and subjected to reduction, alkylation, and in-gel tryptic digestion as described^82,83^. For in-solution analysis, samples were supplemented with 5% SDS, reduced with 10 mM DTT for 15 min at 80 °C, alkylated with 20 mM iodoacetamide for 30 min at RT, then supplemented with a final concentration of 1.2% phosphoric acid and 609 µL of S-Trap (Protifi) binding buffer (90% MeOH/100 mM triethylammonium bicarbonate [TEAB]). Proteins were trapped on an S-Trap micro cartridge, digested using 20 ng/µL sequencing-grade trypsin (Promega) for 1 hr at 47 °C, and eluted using 50 mM TEAB, followed by 0.2% formic acid (FA), followed by 50% acetonitrile (ACN)/0.2% FA. All samples were then lyophilized to dryness. Dried samples were subjected to chromatographic separation on a Waters NanoAquity UPLC equipped with a 1.7 µm BEH130 C18 75 µm I.D. × 250 mm reversed-phase column. The mobile phase consisted of (A) 0.1% FA in water and (B) 0.1% FA in ACN. Following a 4 µL injection, peptides were trapped for 3 min on a 5 µm Symmetry C18 180 µm I.D. × 20 mm column at 5 µL/min in 99.9% A. The analytical column was then switched in-line, and a linear elution gradient of 5% B to 40% B was performed over 60 min at 400 nL/min. The analytical column was connected to a fused silica PicoTip emitter (New Objective) with a 10 µm tip orifice and coupled to a Lumos mass spectrometer (Thermo Scientific) through an electrospray interface operating in data-dependent acquisition mode. The instrument was set to acquire a precursor MS scan from m/z 350 to 1800 every 3 s. In data-dependent mode, MS/MS scans of the most abundant precursors were collected at r=15,000 (45 ms, AGC 5e4) following higher-energy collisional dissociation (HCD) fragmentation at an HCD collision energy of 27%. Within the MS/MS spectra, if any diagnostic O-GlcNAc fragment ions (m/z 204.0867, 138.0545, or 366.1396) were observed, a second MS/MS spectrum at r=30,000 (250 ms, 3e5) of the precursor was acquired with electron transfer dissociation (ETD)/HCD fragmentation using charge-dependent ETD reaction times and either 30% (2+ charge state) or 15% (3+-5+ charge state) supplemental collision energy.

For all experiments, a 60 s dynamic exclusion was employed for previously fragmented precursor ions. Raw liquid chromatography-tandem MS (LC-MS/MS) data files were processed in Proteome Discoverer (Thermo Fisher Scientific) and then submitted to independent Mascot searches (Matrix Science) against a SwissProt database (human taxonomy) containing both forward and reverse entries of each protein (https://www.uniprot.org/proteomes/UP000005640) (20,322 forward entries). Search tolerances were 2 ppm for precursor ions and 0.02 Da for product ions using semi-trypsin specificity with up to two missed cleavages. Both y/b-type HCD and c/z-type ETD fragment ions were allowed for interpreting all spectra. Carbamidomethylation (+57.0214 Da on C) was set as a fixed modification, whereas oxidation (+15.9949 Da on M), phosphorylation (+79.97 Da on S/T) and O-GlcNAc (+203.0794 Da on S/T) were considered dynamic mass modifications. All searched spectra were imported into Scaffold (v4.1, Proteome Software), and scoring thresholds were set to achieve a peptide false discovery rate of 1% using the PeptideProphet algorithm (http://peptideprophet.sourceforge.net/). When satisfactory ETD fragmentation was not obtained upon manual inspection, HCD fragmentation was used to determine O-GlcNAc residue modification using the number of HexNAcs identified in combination with the number of S/T residues in the peptide.

Raw and processed datasets for all proteomics and O-GlcNAc site-mapping experiments are available via ProteomeXchange. [Dataset access details will be added here for reviewers upon journal submission.]

### GlcNDAz crosslinking

GlcNDAz experiments were performed essentially as described^91^. 293T cells were transfected with UAP1^F383G^-myc-6xHis and NF-L-myc-6xHis for 24 hrs, then treated with DMSO or 100 µM Ac_3_GlcNDAz-1P(Ac-SATE)_2_ (“GlcNDAz”) every 24 hrs over 48 hrs. Just before crosslinking, medium was removed and replaced with PBS. With lids removed, culture dishes were placed on ice and underneath a 365 nm UV light (Blak-Ray XX-20BLB UV Bench Lamp, 95-0045-04) for 25 min. Cells were scraped into cold PBS and lysed in 8 M urea/PBS, and lysates were analyzed by IB.

### GlcNDAz crosslink proteomics

293T cells treated with DMSO (vehicle) or GlcNDAz precursor from 30 15 cm dishes were harvested, lysed, and quantified as for IP/co-IP. 77 mg of lysate was divided evenly into 7 5-mL centrifuge tubes and rotated with myc antibody overnight at 4 °C. The next day, 50 µL of settled, washed A/G UltraLink resin was added, rotated at RT for 2 hrs, pooled, and rotated in 600 µL buffer A twice, each 30 min at 4 °C. Ni-NTA purification from the cleared supernatant (1.2 ml total) was performed as above. Resin was washed five times with buffer A and eluted twice with 50 µL elution buffer (250 mM imidazole in 8 M urea/PBS) for 15 min with vigorous shaking. Eluates were separated by SDS-PAGE and stained with colloidal blue. Bands corresponding to NF-L crosslinks were excised by hand, digested in-gel as above, and analyzed by MS/MS proteomics by the Duke Proteomics and Metabolomics Shared Resource.

Raw and processed datasets for all proteomics and O-GlcNAc site-mapping experiments are available via ProteomeXchange. [Dataset access details will be added here for reviewers upon journal submission.]

### Differential extraction

Differential extraction experiments were performed essentially as described^81^. 24 hrs post-transfection, cells were washed three times with 8 mL of 2 mM MgCl_2_/PBS at RT, and rotated in 1 mL of ice-cold low ionic strength (LIS) buffer (10 mM MOPS pH 7, 10 mM MgCl_2_, 1 mM EGTA, 0.15% Triton X-100, protease inhibitor cocktail (1:100) in PBS) for 3 min at RT. The supernatant was collected as the LIS fraction. Remaining cells on the dish were incubated in 1 mL of ice-cold high ionic strength (HIS) buffer (10 mM MOPS pH 7, 10 mM MgCl_2_, 1% Triton X-100, protein inhibitor cocktail (1:100) and Benzonase nuclease (Novagen, 70746, 1:100) in PBS) for 3 min on ice. Then, 250 µL of ice-cold 5 M NaCl (final 1 M NaCl) was added, cells were resuspended by pipetting and transferred to a clean microfuge tube as the HIS fraction. LIS and HIS fractions were centrifuged at 27,000 g for 15 min at 4 °C, and cleared supernatants were exchanged into IP lysis buffer with 0.1% SDS using Zeba spin columns. The insoluble pellets from both LIS and HIS tubes were extracted in 200 µL of 8 M urea/PBS on ice for 20 min and probe-sonicated for 50 s at 50% duty cycle. Supernatant was then exchanged to the IP lysis buffer with 0.1% SDS buffer and labeled as the insoluble fraction.

### IFA

100,000 *NEFL*^-/-^ SH-SY5Y cells seeded in 12-well plates with an 18 mm coverslip on the bottom were transfected with 0.2 µg of each DNA (e.g., NF-L, OGT). 24 hrs later, the medium was changed to fresh DMEM/F12. 48 hrs after transfection, cells were washed twice with PBS, fixed with 4% paraformaldehyde (PFA, MP Biomedicals, 02150146.5 diluted in water) at RT for 15 min, permeabilized with 0.1% Triton X-100/PBS at RT for 10 min, and incubated in blocking buffer (1% BSA/PBS) at RT for 1 hr. Coverslips were incubated with the RL2 and NF-L antibodies (1:400 in blocking buffer) overnight at 4 °C, washed three times with PBS and incubated in Alexa Fluor-conjugated secondary antibody (1:400 in blocking buffer) in the dark at RT for 1 hr. Coverslips were washed with PBS twice before mounting in ProLong Diamond anti-fade mounting medium with DAPI (Invitrogen, P36931) onto the microscope slides. For NF-L/INA co-expression, 200,000 SW13 vim^-^ cells seeded in a 12-well plate were transfected with 0.1 µg of each DNA using Lipofectamine 3000, processed as above and stained with the myc and V5 antibodies (1:400 in blocking buffer). Quantification of NF-L morphology in 5-7 cells per condition per replicate was performed by a blinded investigator. For rat hippocampal neurons, cells were fixed for 8 min using warm 4% PFA/4% sucrose, permeabilized with 0.1% Triton X-100/PBS at RT for 5 min, blocked for 1 hr in blocking solution (5% goat serum and 1% BSA/PBS), and stained with Tuj1 and mCherry antibodies (1:400 in blocking buffer). Following three PBS washes after secondary antibodies, cells were incubated with Hoechst 33342 reagent (ThermoFisher, H3570, 1:2,000 in blocking buffer) for 10 min and then washed once with PBS prior to image acquisition. Complete information for all antibodies is provided in Table 3.

### Image acquisition

Cells were imaged on an inverted Zeiss 780 single point scanning confocal microscope equipped with a fully motorized Zeiss Axio Observer microscope base, Marzhauser linearly encoded stage, diode (405 nm), argon ion (488 nm), double solid-state (561 nm), and helium-neon (633 nm) lasers. Images were acquired at RT using a 63x NA/1.4 oil plan apochromatic oil immersion objective lens. Images were acquired sequentially by frame scanning bidirectionally using the galvanometer-based imaging mode in Zeiss Zen Black Acquisition software and processed using Fiji ImageJ. Detection ranges were 480-500 nm, 490-550 nm, and 650-750 nm.

### Live-cell imaging

On day 6 *in vitro*, neurons were transfected with 0.3 µg of each DNA using Lipofectamine 3000. 24 hrs later, 5-7 axons per condition per replicate were imaged live every 5 s over 5 min at 10 ms exposure on a Zeiss ELRYA7 super-resolution microscope equipped with a Pecon environmental chamber at 37 °C, lattice SIM, and Plan-Apochromat 63x/1.4 Oil DIC M27 lens. Time lapse plus Z imaging was performed with laser excitation at 405 nm, 488 nm, z-step size 110 nm, SIM grating size of 27 µm, and number of SIM phases (13). Laser power was set to < 1% for each channel to minimize phototoxicity during acquisition. Raw SIM data were processed using SIM Processing in the Zen Software (Zeiss), with “adjusted” setting “normal to baseline cut,” followed by “maximum intensity projection.” Then, data were processed using IMARIS software with the “surface” function and “track over time” to track distance, displacement, and speed of mitochondria and lysosomes as described^85^.

## Statistical analysis

All experimental data presented are representative of at least three independent biological replicates unless otherwise indicated. All statistical analyses were performed in Prism 9 using one-way ANOVA followed by Tukey’s post-hoc correction, two-tailed students test, or Mann-Whitney test, followed by Dunn’s correction. n.s,. not significant; *, p < 0.05; **, p < 0.005;

***, p < 0.0005; ****, p < 0.0001.

## References

1 Hohmann, T. & Dehghani, F. The Cytoskeleton-A Complex Interacting Meshwork. Cells 8, doi:10.3390/cells8040362 (2019).

2 Yuan, A., Rao, M. V., Veeranna & Nixon, R. A. Neurofilaments and Neurofilament Proteins in Health and Disease. Cold Spring Harb Perspect Biol. 9, a018309, doi:10.1101/cshperspect.a018309 (2017).

3 Yuan, A. & Nixon, R. A. Neurofilament Proteins as Biomarkers to Monitor Neurological Diseases and the Efficacy of Therapies. Front Neurosci 15, 689938, doi:10.3389/fnins.2021.689938 (2021).

4 Gafson, A. R. et al. Neurofilaments: neurobiological foundations for biomarker applications. Brain 143, 1975–1998, doi:10.1093/brain/awaa098 (2020).

5 Herrmann, H. & Aebi, U. Intermediate Filaments: Structure and Assembly. Cold Spring Harb Perspect Biol 8, a018242, doi:10.1101/cshperspect.a018242 (2016).

6 Lowery, J., Kuczmarski, E. R., Herrmann, H. & Goldman, R. D. Intermediate Filaments Play a Pivotal Role in Regulating Cell Architecture and Function. J Biol Chem. 290, 17145–17153, doi:10.1074/jbc.R115.640359 (2015).

7 Bott, C. J. W. B. Intermediate filaments in developing neurons: Beyond structure. Cytoskeleton, 110–128, doi:10.1002/cm.21597 (2020).

8 van Bodegraven, E. J. & Etienne-Manneville, S. Intermediate Filaments from Tissue Integrity to Single Molecule Mechanics. Cells 10, doi:10.3390/cells10081905 (2021).

9 Chernyatina, A. A., Hess, J. F., Guzenko, D., Voss, J. C. & Strelkov, S. V. How to Study Intermediate Filaments in Atomic Details. Methods Enzymol. 568, 3–33, doi:10.1016/bs.mie.2015.09.024 (2016).

10 Köster, S., Weitz, D. A., Goldman, R. D., Aebi, U. & Herrmann, H. Intermediate filament mechanics in vitro and in the cell: from coiled coils to filaments, fibers and networks. Curr Opin Cell Biol 32, 82–91, doi:10.1016/j.ceb.2015.01.001 (2015).

11 Leduc, C. & Etienne-Manneville, S. Intermediate filaments in cell migration and invasion: the unusual suspects. Curr Opin Cell Biol 32, 102–112, doi:10.1016/j.ceb.2015.01.005 (2015).

12 Kreplak, L., Bär, H., Leterrier, J. F., Herrmann, H. & Aebi, U. Exploring the mechanical behavior of single intermediate filaments. J Mol Biol 354, 569–577, doi:10.1016/j.jmb.2005.09.092 (2005).

13 Zhu, Q., Couillard-Després, S. & Julien, J. P. Delayed maturation of regenerating myelinated axons in mice lacking neurofilaments. Exp Neurol. 148, 299–316, doi:10.1006/exnr.1997.6654. (1997).

14 Zhang, Z., Casey, D. M., Julien, J.-P. & Xu, Z. Normal dendritic arborization in spinal motoneurons requires neurofilament subunit L. J Comp Neurol. 450, 144–152, doi:10.1002/cne.10306 (2002).

15 Blizzard, C. A., King, A. E., Vickers, J. & Dickson, T. Cortical Murine Neurons Lacking the Neurofilament Light Chain Protein Have an Attenuated Response to Injury In Vitro. J Neurotrauma. 30, 1908–1918, doi:10.1089/neu.2013.2850 (2013).

16 Sainio, M. T. et al. Neurofilament Light Regulates Axon Caliber, Synaptic Activity, and Organelle Trafficking in Cultured Human Motor Neurons. Front Cell Dev Biol. 9, 820105, doi:10.3389/fcell.2021.820105 (2022).

17 Yuan, A. et al. Neurofilament subunits are integral components of synapses and modulate neurotransmission and behavior in vivo. Mol Psychiatry. 20, 986–994, doi:10.1038/mp.2015.45 (2015).

18 Ehlers, M. D., Fung, E. T., O’Brien, R. J. & Huganir, R. L. Splice variant-specific interaction of the NMDA receptor subunit NR1 with neuronal intermediate filaments. J Neurosci. 18, 720–730, doi:10.1523/JNEUROSCI.18-02-00720.1998 (1998).

19 Yuan, A. et al. Neurofilament light interaction with GluN1 modulates neurotransmission and schizophrenia-associated behaviors. Transl Psychiatry 8, 167, doi:10.1038/s41398-018-0194-7 (2018).

20 Imarisio, A. et al. Synaptic neurofilaments and GluN1-NfL interaction in experimental models of α-synucleinopathies. Neurodegener Dis, doi:10.1159/000526376 (2022).

21 Stone, E. J., Kolb, S. J. & Brown, A. A review and analysis of the clinical literature on Charcot-Marie-Tooth disease caused by mutations in neurofilament protein L. Cytoskeleton (Hoboken) 78, 97–110, doi:10.1002/cm.21676 (2021).

22 Kim, H. J. et al. Phenotypic heterogeneity in patients with NEFL-related Charcot-Marie-Tooth disease. Mol Genet Genomic Med., e1870, doi:10.1002/mgg3.1870 (2022).

23 Laurá, M., Pipis, M., Rossor, A. M. & Reilly, M. M. Charcot-Marie-Tooth disease and related disorders: an evolving landscape. Curr Opin Neurol. 32, 641–650, doi:10.1097/WCO.0000000000000735 (2019).

24 Gentil, B. J. et al. Normal role of the low-molecular-weight neurofilament protein in mitochondrial dynamics and disruption in Charcot-Marie-Tooth disease. FASEB J. 26, 1194–1203, doi:10.1096/fj.11-196345 (2012).

25 Israeli, E. et al. Intermediate filament aggregates cause mitochondrial dysmotility and increase energy demands in giant axonal neuropathy. Hum Mol Genet. 24, 2143–2157, doi:10.1093/hmg/ddw081 (2016).

26 Van Lent, J. et al. Induced pluripotent stem cell-derived motor neurons of CMT type 2 patients reveal progressive mitochondrial dysfunction. Brain 144, 2471–2485, doi:10.1093/brain/awab226 (2021).

27 Adebola, A. A. et al. Neurofilament light polypeptide gene N98S mutation in mice leads to neurofilament network abnormalities and a Charcot-Marie-Tooth Type 2E phenotype. Hum Mol Genet 24, 2163–2174, doi:10.1093/hmg/ddu736 (2015).

28 Saporta, M. A. et al. Axonal Charcot-Marie-Tooth Disease Patient-Derived Motor Neurons Demonstrate Disease-Specific Phenotypes Including Abnormal Electrophysiological Properties. Exp Neurol 263, 190–199, doi:10.1016/j.expneurol.2014.10.005 (2015).

29 Brownlees, J. et al. Charcot-Marie-Tooth disease neurofilament mutations disrupt neurofilament assembly and axonal transport. Hum Mol Genet. 11, 2837–2844, doi:10.1093/hmg/11.23.2837. (2002).

30 Zhao, J., Brown, K. & Liem, R. K. H. Abnormal neurofilament inclusions and segregations in dorsal root ganglia of a Charcot-Marie-Tooth type 2E mouse model. PLoS One 12, e0180038, doi:10.1371/journal.pone.0180038 (2017).

31 Ishii, T., Haga, S. & Tokutake, S. Presence of neurofilament protein in Alzheimer’s neurofibrillary tangles (ANT). An immunofluorescent study. Acta Neuropathol 48, 105–112, doi:10.1007/bf00691151 (1979).

32 Goldman, J. E., Yen, S. H., Chiu, F. C. & Peress, N. S. Lewy bodies of Parkinson’s disease contain neurofilament antigens. Science 221, 1082–1084, doi:10.1126/science.6308771 (1983).

33 Delisle, M. B. & Carpenter, S. Neurofibrillary axonal swellings and amyotrophic lateral sclerosis. J Neurol Sci 63, 241–250, doi:10.1016/0022-510x(84)90199-0 (1984).

34 Franco-Espin, J. et al. SMN Is Physiologically Downregulated at Wild-Type Motor Nerve Terminals but Aggregates Together with Neurofilaments in SMA Mouse Models. Biomolecules 12, doi:10.3390/biom12101524 (2022).

35 Khalil, M. et al. Neurofilaments as biomarkers in neurological disorders. Nat Rev Neurol. 14, 577–589, doi:10.1038/s41582-018-0058-z (2018).

36 Chelban, V. et al. Neurofilament light levels predict clinical progression and death in multiple system atrophy. Brain 145, 4398–4408, doi:10.1093/brain/awac253 (2022).

37 Fox, R. J. et al. Neurofilament light chain in a phase 2 clinical trial of ibudilast in progressive multiple sclerosis. Mult Scler 27, 2014–2022, doi:10.1177/1352458520986956 (2021).

38 Rival, M. et al. Neurofilament Light Chain Levels Are Predictive of Clinical Conversion in Radiologically Isolated Syndrome. Neurol Neuroimmunol Neuroinflamm 10, doi:10.1212/nxi.0000000000200044 (2023).

39 Thebault, S. et al. Serum neurofilament light chain predicts long term clinical outcomes in multiple sclerosis. Sci Rep 10, 10381, doi:10.1038/s41598-020-67504-6 (2020).

40 Bos, I. et al. Cerebrospinal fluid biomarkers of neurodegeneration, synaptic integrity, and astroglial activation across the clinical Alzheimer’s disease spectrum. Alzheimers Dement 15, 644–654, doi:10.1016/j.jalz.2019.01.004 (2019).

41 Snider, N. T. & Omary, M. B. Post-translational modifications of intermediate filament proteins: mechanisms and functions. Nat Rev Mol Cell Biol. 15, 163–177, doi:10.1038/nrm3753 (2014).

42 Hisanaga, S., Gonda, Y., Inagaki, M., Ikai, A. & Hirokawa, N. Effects of phosphorylation of the neurofilament L protein on filamentous structures. Cell Regul. 1, 237–248, doi:10.1091/mbc.1.2.237 (1990).

43 Gonda, Y. et al. Involvement of protein kinase C in the regulation of assembly-disassembly of neurofilaments in vitro. Biochem Biophys Res Commun. 167, 1316–1325, doi:10.1016/0006-291x(90)90667-c (1990).

44 Yates, D. M. et al. Neurofilament subunit (NFL) head domain phosphorylation regulates axonal transport of neurofilaments. Eur J Cell Biol. 88, 193–202, doi:10.1016/j.ejcb.2008.11.004 (2009).

45 Veeranna et al. Mitogen-activated protein kinases (Erk1,2) phosphorylate Lys-Ser-Pro (KSP) repeats in neurofilament proteins NF-H and NF-M. J Neurosci. 18, 4008–4021, doi:10.1523/JNEUROSCI.18-11-04008.1998 (1998).

46 Kesavapany, S., Li, B. S. & Pant, H. C. Cyclin-dependent kinase 5 in neurofilament function and regulation. Neurosignals 12, 252–264, doi:10.1159/000074627. (2003).

47 Dong, D. L. et al. Glycosylation of Mammalian Neurofilaments. Localization of Multiple O-linked N-acetylglucosamine Moieties on Neurofilament Polypeptides L and M. J Biol Chem. 268, 16679–16687 (1993).

48 Dong, D. L., Xu, Z. S., Hart, G. W. & Cleveland, D. W. Cytoplasmic O-GlcNAc modification of the head domain and the KSP repeat motif of the neurofilament protein neurofilament-H. J Biol Chem. 271, 20845–20852, doi:10.1074/jbc.271.34.20845 (1996).

49 Vosseller, K. et al. O-linked N-acetylglucosamine proteomics of postsynaptic density preparations using lectin weak affinity chromatography and mass spectrometry. Mol Cell Proteomics 5, 923–934, doi:10.1074/mcp.T500040-MCP200 (2006).

50 Trinidad, J. C. et al. Global identification and characterization of both O-GlcNAcylation and phosphorylation at the murine synapse. Mol Cell Proteomics 11, 215–229, doi:10.1074/mcp.O112.018366 (2012).

51 Lee, B. E. et al. O-GlcNAcylation regulates dopamine neuron function, survival and degeneration in Parkinson disease. Brain 143, 3699–3716, doi:10.1093/brain/awaa320 (2020).

52 Wang, S. et al. Quantitative Proteomics Identifies Altered O-GlcNAcylation of Structural, Synaptic and Memory-Associated Proteins in Alzheimer’s Disease. J Pathol. 243, 78–88, doi:10.1002/path.4929 (2017).

53 Zachara, N. E., Akimoto, Y., Boyce, M. & Hart, G. W. in Essentials of Glycobiology (eds th et al.) 251–264 (2022).

54 Bond, M. R. & Hanover, J. A. A little sugar goes a long way: the cell biology of O-GlcNAc. J Cell Biol. 208, 869–880, doi:10.1083/jcb.201501101 (2015).

55 Hart, G. W. Three Decades of Research on O-GlcNAcylation - A Major Nutrient Sensor That Regulates Signaling, Transcription and Cellular Metabolism. Front Endocrinol (Lausanne) 5, 183, doi:10.3389/fendo.2014.00183 (2014).

56 Keembiyehetty, C. et al. Conditional knock-out reveals a requirement for O-linked N-Acetylglucosaminase (O-GlcNAcase) in metabolic homeostasis. J Biol Chem. 290, 7097–7113, doi:10.1074/jbc.M114.617779 (2015).

57 Yang, Y. R. et al. O-GlcNAcase is essential for embryonic development and maintenance of genomic stability. Aging Cell 11, 439–448, doi:10.1111/j.1474-9726.2012.00801.x (2012).

58 Akimoto, Y. et al. Localization of the O-GlcNAc transferase and O-GlcNAc-modified proteins in rat cerebellar cortex. Brain Res. 966, 194–205, doi:10.1016/s0006-8993(02)04158-6 (2003).

59 Cole, R. N. & Hart, G. W. Cytosolic O-glycosylation is abundant in nerve terminals. J Neurochem. 79, 1080–1089, doi:10.1046/j.1471-4159.2001.00655.x (2001).

60 Lagerlöf, O. et al. The nutrient sensor OGT in PVN neurons regulates feeding. Science. 351, 1293–1296, doi:10.1126/science.aad5494 (2016).

61 Wang, Q. et al. Ventromedial hypothalamic OGT drives adipose tissue lipolysis and curbs obesity. Sci Adv 8, eabn8092, doi:10.1126/sciadv.abn8092 (2022).

62 Huynh, D. T. & Boyce, M. Chemical Biology Approaches to Understanding Neuronal O−GlcNAcylation. Israel J Chem., doi:10.1002/ijch.202200071 (2022).

63 Fehl, C., Hanover, J.A. Tools, tactics and objectives to interrogate cellular roles of O-GlcNAc in disease. Nat Chem Biol., doi:10.1038/s41589-021-00903-6 (2021).

64 Liu, F., Iqbal, K., Grundke-Iqbal, I., Hart, G. W. & Gong, C.-X. O-GlcNAcylation regulates phosphorylation of tau: A mechanism involved in Alzheimer’s disease. Proc Natl Acad Sci U S A. 101, 10804–10809, doi:10.1073/pnas.0400348101 (2004).

65 Marotta, N. P. et al. O-GlcNAc modification blocks the aggregation and toxicity of the protein α-synuclein associated with Parkinson’s disease. Nature Chem. 7, 913–920, doi:10.1038/nchem.2361 (2015).

66 Park, J., Lai, M. K. P., Arumugam, T. V. & Jo, D. G. O-GlcNAcylation as a Therapeutic Target for Alzheimer’s Disease. Neuromolecular Med. 22, 171–193, doi:10.1007/s12017-019-08584-0 (2020).

67 Balana, A. T. & Pratt, M. R. Mechanistic roles for altered O-GlcNAcylation in neurodegenerative disorders. Biochem J. 478, 2733–2758, doi:10.1042/BCJ20200609 (2021).

68 Permanne, B. et al. O-GlcNAcase Inhibitor ASN90 is a Multimodal Drug Candidate for Tau and α-Synuclein Proteinopathies. ACS Chem Neurosci. 13, 1296–1314, doi:10.1021/acschemneuro.2c00057. (2022).

69 Yuzwa, S. A. et al. A potent mechanism-inspired O-GlcNAcase inhibitor that blocks phosphorylation of tau in vivo. Nat Chem Biol. 4, 483–490, doi:10.1038/nchembio.96 (2008).

70 Yuzwa, S. A. et al. Increasing O-GlcNAc slows neurodegeneration and stabilizes tau against aggregation. Nat Chem Biol. 8, 393–399, doi:10.1038/nchembio.797 (2012).

71 Wang, X. et al. Early intervention of tau pathology prevents behavioral changes in the rTg4510 mouse model of tauopathy. PLoS One 13, e0195486, doi:10.1371/journal.pone.0195486 (2018).

72 Wang, X. et al. MK-8719, a Novel and Selective O-GlcNAcase Inhibitor That Reduces the Formation of Pathological Tau and Ameliorates Neurodegeneration in a Mouse Model of Tauopathy. J Pharmacol Exp Ther. 374, 252–263, doi:10.1124/jpet.120.266122. (2020).

73 Kielbasa, W. et al. A single ascending dose study in healthy volunteers to assess the safety and PK of LY3372689, an inhibitor of O-GlcNAcase (OGA) enzyme. Alzheimers Dement., doi:10.1002/alz.040473 (2020).

74 Sandhu, P. et al. in Alzheimers’ Association International Conference (AAIC):P4 (2016).

75 Lowe, S. L. et al. Single and multiple ascending dose studies in healthy volunteers to assess the safety and PK of LY3372689, an inhibitor of the O-GlcNAcase (OGA) enzyme. Alzheimers Dement., doi:10.1002/alz.057728 (2021).

76 Lazarus, M. B., Nam, Y., Jiang, J., Sliz, P. & Walker, S. Structure of human O-GlcNAc transferase and its complex with a peptide substrate. Nature 469, 564–567, doi:10.1038/nature09638 (2011).

77 Gloster, T. M. et al. Hijacking a biosynthetic pathway yields a glycosyltransferase inhibitor within cells. Nat Chem Biol. 7, 174–181, doi:10.1038/nchembio.520 (2011).

78 Zachara, N. E., Cole, R. N., Hart, G. W. & Gao, Y. Detection and analysis of proteins modified by O-linked N-acetylglucosamine. Curr Protoc Protein Sci Chapter 12, Unit 12.18, doi:10.1002/0471140864.ps1208s25 (2001).

79 Dalvai, M. et al. Scalable Genome-Editing-Based Approach for Mapping Multiprotein Complexes in Human Cells. Cell Rep. 13, 621–633, doi:10.1016/j.celrep.2015.09.009 (2015).

80 Myers, S. A., Daou, S., Affar el, B. & Burlingame, A. Electron transfer dissociation (ETD): the mass spectrometric breakthrough essential for O-GlcNAc protein site assignments-a study of the O-GlcNAcylated protein host cell factor C1. Proteomics 13, 982–991, doi:10.1002/pmic.201200332 (2013).

81 Tarbet, H. J. et al. Site-specific glycosylation regulates the form and function of the intermediate filament cytoskeleton. eLife 7, e31807, doi:10.7554/eLife.31807 (2018).

82 Chen, P. H. et al. Gigaxonin glycosylation regulates intermediate filament turnover and may impact giant axonal neuropathy etiology or treatment. JCI Insight 5, e127751, doi:10.1172/jci.insight.127751 (2019).

83 Bisnett, B. J. et al. Evidence for nutrient-dependent regulation of the COPII coat by O-GlcNAcylation. Glycobiology 31, 1102–1120, doi:10.1093/glycob/cwab055 (2021).

84 Ridge, K. M. et al. Methods for Determining the Cellular Functions of Vimentin Intermediate Filaments. Methods Enzymol 568, 389–426, doi:10.1016/bs.mie.2015.09.036 (2016).

85 Evans, C. S. & Holzbaur, E. L. Degradation of engulfed mitochondria is rate-limiting in Optineurin-mediated mitophagy in neurons. Elife 9, doi:10.7554/eLife.50260 (2020).

86 Bocquet, A. et al. Neurofilaments bind tubulin and modulate its polymerization. J Neurosci 29, 11043–11054, doi:10.1523/jneurosci.1924-09.2009 (2009).

87 Yadav, P. et al. Neurofilament depletion improves microtubule dynamics via modulation of Stat3/stathmin signaling. Acta Neuropathologica 132, 93–110, doi:10.1007/s00401-016-1564-y (2016).

88 Cason, S. E. & Holzbaur, E. L. F. Selective motor activation in organelle transport along axons. Nat Rev Mol Cell Biol 23, 699–714, doi:10.1038/s41580-022-00491-w (2022).

89 Tarbet, H. J., Toleman, C. A. & Boyce, M. A Sweet Embrace: Control of Protein-Protein Interactions by O-Linked β-N-Acetylglucosamine. Biochemistry 57, 13–21, doi:10.1021/acs.biochem.7b00871 (2018).

90 Carter, J. et al. Neurofilament (NF) assembly; divergent characteristics of human and rodent NF-L subunits. J Biol Chem. 273, 5101–5108, doi:10.1074/jbc.273.9.5101 (1998).

91 Yu, S.-H. et al. Metabolic labeling enables selective photocrosslinking of O-GlcNAc-modified proteins to their binding partner. Proc Natl Acad Sci U S A. 109, 4834–4839, doi:10.1073/pnas.1114356109 (2012).

92 Yuan, A. et al. Alpha-internexin Is Structurally and Functionally Associated With the Neurofilament Triplet Proteins in the Mature CNS. J Neurosci. 26, 10006–10019, doi:10.1523/JNEUROSCI.2580-06.2006 (2006).

93 Cairns, N. J. et al. Alpha-internexin is present in the pathological inclusions of neuronal intermediate filament inclusion disease. Am J Pathol 164, 2153–2161, doi:10.1016/s0002-9440(10)63773-x (2004).

94 Zhao, J. & Liem, R. K. Alpha-Internexin and Peripherin: Expression, Assembly, Functions, and Roles in Disease. Methods Enzymol 568, 477–507, doi:10.1016/bs.mie.2015.09.012 (2016).

95 Sarria, A. J., Lieber, J. G., Nordeen, S. K. & Evans, R. M. The presence or absence of a vimentin-type intermediate filament network affects the shape of the nucleus in human SW-13 cells. J Cell Sci 107 (Pt 6), 1593–1607, doi:10.1242/jcs.107.6.1593 (1994).

96 Ku, N. O., Toivola, D. M., Strnad, P. & Omary, M. B. Cytoskeletal keratin glycosylation protects from epithelial tissue injury. Nat Cell Biol. 12, 876–885, doi:10.1038/ncb2091 (2010).

97 Srikanth, B., Vaidya, M. M. & Kalraiya, R. D. O-GlcNAcylation determines the solubility, filament organization, and stability of keratins 8 and 18. J Biol Chem. 285 34062–34071, doi:10.1074/jbc.M109.098996 (2010).

98 Sasaki, T. et al. Aggregate formation and phosphorylation of neurofilament-L Pro22 Charcot-Marie-Tooth disease mutants. Hum Mol Genet. 15, 943–952, doi:10.1093/hmg/ddl011. (2006).

99 Liao, L. et al. Proteomic characterization of postmortem amyloid plaques isolated by laser capture microdissection. J Biol Chem 279, 37061–37068, doi:10.1074/jbc.M403672200 (2004).

100 Manetto, V., Sternberger, N. H., Perry, G., Sternberger, L. A. & Gambetti, P. Phosphorylation of neurofilaments is altered in amyotrophic lateral sclerosis. J Neuropathol Exp Neurol. 47, 642–653, doi:10.1097/00005072-198811000-00007 (1988).

101 Hart, G. W., Slawson, C., Ramirez-Correa, G. & Lagerlof, O. Cross talk between O-GlcNAcylation and phosphorylation: roles in signaling, transcription, and chronic disease. Annu Rev Biochem. 80, 825–858, doi:10.1146/annurev-biochem-060608-102511 (2011).

102 Wang, S. et al. Extensive crosstalk between O-GlcNAcylation and phosphorylation regulates Akt signaling. PLoS ONE 7, e37427, doi:10.1371/journal.pone.0037427 (2012).

103 Trinidad, J. C. et al. Global identification and characterization of both O-GlcNAcylation and phosphorylation at the murine synapse. Mol Cell Proteomics 11, 215–229, doi:10.1074/mcp.O112.018366 (2012).

104 Zhong, J. et al. Quantitative phosphoproteomics reveals crosstalk between phosphorylation and O-GlcNAc in the DNA damage response pathway. Proteomics 15, 591–607, doi:10.1002/pmic.201400339 (2015).

105 Leney, A. C., El Atmioui, D., Wu, W., Ovaa, H. & Heck, A. J. R. Elucidating crosstalk mechanisms between phosphorylation and O-GlcNAcylation. Proc Natl Acad Sci U S A 114, E7255–e7261, doi:10.1073/pnas.1620529114 (2017).

106 Toleman, C. A. et al. Structural basis of O-GlcNAc recognition by mammalian 14-3-3 proteins. Proc Natl Acad Sci U S A 115, 5956–5961, doi:10.1073/pnas.1722437115 (2018).

107 Cleverley, K. E., Betts, J. C., Blackstock, W. P., Gallo, J. M. & Anderton, B. H. Identification of novel in vitro PKA phosphorylation sites on the low and middle molecular mass neurofilament subunits by mass spectrometry. Biochemistry 37, 3917–3930, doi:10.1021/bi9724523 (1998).

108 Trinidad, J. C., Specht, C. G., Thalhammer, A., Schoepfer, R. & Burlingame, A. L. Comprehensive identification of phosphorylation sites in postsynaptic density preparations. Mol Cell Proteomics 5, 914–922, doi:10.1074/mcp.T500041-MCP200 (2006).

109 Lundby, A. et al. Quantitative maps of protein phosphorylation sites across 14 different rat organs and tissues. Nat Commun 3, 876, doi:10.1038/ncomms1871 (2012).

110 Sacco, F. et al. Glucose-regulated and drug-perturbed phosphoproteome reveals molecular mechanisms controlling insulin secretion. Nat Commun 7, 13250, doi:10.1038/ncomms13250 (2016).

111 Huynh, V. N. et al. Defining the Dynamic Regulation of O-GlcNAc Proteome in the Mouse Cortex---the O-GlcNAcylation of Synaptic and Trafficking Proteins Related to Neurodegenerative Diseases. Front Aging 2, 757801, doi:10.3389/fragi.2021.757801 (2021).

112 Alfaro, J. F. et al. Tandem mass spectrometry identifies many mouse brain O-GlcNAcylated proteins including EGF domain-specific O-GlcNAc transferase targets. Proc Nat Acad Sci U S A 109, 7280–7285, doi:10.1073/pnas.1200425109 (2012).

113 Eldirany, S. A., Lomakin, I. B., Ho, M. & Bunick, C. G. Recent insight into intermediate filament structure. Curr Opin Cell Biol. 68, 132–143, doi:doi.org/10.1016/j.ceb.2020.10.001 (2021).

114 Cheung, W. D. & Hart, G. W. AMP-activated Protein Kinase and p38 MAPK Activate O-GlcNAcylation of Neuronal Proteins during Glucose Deprivation. J Biol Chem. 283, 13009–13020 (2008).

115 Pekkurnaz, G., Trinidad, J. C., Wang, X., Kong, D. & Schwarz, T. L. Glucose regulates mitochondrial motility via Milton modification by O-GlcNAc transferase. Cell 158, 54–68, doi:10.1016/j.cell.2014.06.007 (2014).

116 Zhou, X. et al. Mutations linked to neurological disease enhance self-association of low-complexity protein sequences. Science 377, eabn5582, doi:10.1126/science.abn5582 (2022).

